# Trinucleotide Distribution, Symmetry Elements and Formulation of Mirror Symmetry Index for G4 Motifs

**DOI:** 10.64898/2026.07.05.736592

**Authors:** Arjun Arya, Bhaskar Datta

**Author notes:** **Arjun Arya**.

## Abstract

Symmetry elements in nucleic acids are most strongly correlated with sites of biological function; however, their relevance to non-canonical structures remains underexplored. In this study, we demonstrate the presence and significance of trinucleotide symmetry elements within G-quadruplex (G4) motifs. Our central hypothesis is that the intra-strand mirror symmetry of trinucleotides has been evolutionarily selected to facilitate G4 formation builds on the established sequence-structure association of G-quadruplexes and the natural symmetry law governing nucleotide insertion during genome evolution. Using a conserved G4 motif in the first exon of the MTOR gene as a model, we showed remarkable trinucleotide symmetry preservation across primates and broader mammals, with functional G4 regions displaying locally elevated symmetry relative to the codon-biased exonic background. Analysis of experimentally validated oncogenic G4s, including c-MYC, BCL2, VEGF, and KRAS, revealed that mirror and reverse complement symmetries converge around biologically important G4s. To quantify this feature, we formulated two complementary descriptors: the mirror symmetry index (MSI) and its non-palindromic variant (nMSI). Across 14 oncogene-promoter wild-type G4s, the majority scored MSI ≥ 0.80 (mean 0.884), with only the loop-rich ATG7, BCR, and MDM2 motifs falling below this value, and the KRAS promoter G4 reached individual significance against its mononucleotide-preserving null distribution (p = 0.042). Most decisively, each wild-type G4 scored higher on MSI than its experimentally confirmed G4-abolished mutant in 12 of 14 paired comparisons (sign test, p = 0.0065; mean ΔMSI = +0.089, mean ΔnMSI = +0.192); the two reversals (BCL2 and HIF-1α) are attributable to scrambled mutant controls that introduce more balanced trinucleotide compositions rather than to failure of the index. The directional trend was reproduced across three independently published datasets, with nMSI ≥ 0.50 separating G4-forming from non-G4 sequences at 77.8% sensitivity and 100% specificity, although the collective per-sequence signal from mononucleotide-preserving shuffles remained a non-significant trend (Stouffer combined Z = 1.197, p = 0.116). This first report of trinucleotide symmetry in G4 motifs posits that coordinated nucleotide insertion and quadruplet maintenance act as an evolutionary forcing mechanism that pre-organizes single strands for G4 folding.

## Introduction

The structural symmetries within nucleic acids have long been associated with their energetic stability and biological function. These correlations are inspired by Noether’s demonstrations of mathematical symmetries, which are intricately linked to the fundamental conservation laws in physics[1]. Jacques Monod stressed that a high degree of order and complexity in biological systems appears to be symmetry-aligned albeit without straightforward geometric interpretations[2]. The high degree of order and symmetry in nucleic acids can be extrapolated from the definitive positions of nucleotides in DNA and Chargaff’s second parity rule (CSPR)[3], [4]. Chargaff’s first rule of parity regarding A=T and C=G between two strands of DNA can be completely accounted for by Watson-Crick base pairing CSPR suggests the equality of nucleotide and oligonucleotide frequencies within a single DNA strand[5]. The etiology of this intra-strand symmetry remains intriguing, as it cannot be attributed to inter-strand pairing alone[6], [7]. Recent studies suggest that a natural law of symmetry underlying the creation and preservation of DNA may explain how DNA retains its unique identity[3]. This law assumes that during the evolution of genomes, nucleotides are inserted on the two strands at the same time in a direct reverse (5’ → 3’) sequence[3], [8]. Such a system ensures that mutations or insertions are compensated to keep the genome intact, thereby limiting the entropic gain due to random mutations[3]. The corresponding symmetry forms a quadruplet-based classification system that groups all 64 trinucleotides into A+T-rich and C+G-rich groups[3], [9]. Each quadruplet consists of four types of trinucleotides: direct (D), Reverse Complement (RC), complement (C), and reverse (R). In these quadruplets, a “butterfly” symmetry emerges, with the frequencies of D = RC and C = R being reflected in a single strand[3], [9]. These quadruplets form the fundamental units of the genome; their internal symmetry (D=RC, C=R) differs between species and represents an invariant molecular mark of evolutionary stability[3], [4].

While coding DNA often breaks CSPR, these laws of symmetry are universal in non-coding DNA[4], [10]. Coding sequences are frequently associated with pronounced deviations in the human genome, such as possessing excess adenine over thymine[11]. Such violations are mostly explained by codon usage bias, as the need to have certain amino acids can induce local breakdown of symmetry[12]. Nevertheless, the genome maintains an overall balance through inverse proportionality; local exon asymmetries are replaced by inverse changes in the frequencies of flanking non-coding regions, such that the total nucleotide count and overall genomic symmetry are maintained[4], [13]. A cycle of protection can be inferred that enables the development of species-specific coding characteristics while retaining the overall structural stability of DNA molecule[3].

Within this genetic symmetry, G-rich genomic regions represent distinct loci for the distribution analysis. These regions are also important because they can fold into G-quadruplex (G4) structures[14]. G4s are non-canonical three-dimensional structures stabilized by Hoogsteen hydrogen bonds and stacked guanine tetrads[15], [16], [17], [18]. While G4s were formerly treated as structural curiosities, they are now recognized as omnipresent multi-functional epigenetic regulating elements in promoters, exons, and untranslated regions[16], [19], [20]. G4 motifs have garnered significant interest because of their regulatory functions in important genomic regions, particularly in promoters, 5′ and 3′ untranslated regions (UTRs), telomeres, and the first exons of most genes[18], [21]. Genome-wide analyses have indicated that nearly half of all human gene promoters contain putative G4-forming sequences, which are particularly abundant in oncogenes and transcription regulators[18], [22]. Comparative genomics has demonstrated that G4 motifs, particularly those with shorter loop sizes or conserved sequence contexts, are conserved across species[23], [24], [25], [26], [27], [28]. These conservation patterns align with selective pressure, highlighting their utilitarian functions[16], [18], [29], [30].

We hypothesized that trinucleotide symmetry has been evolutionarily selected to facilitate the formation of G4 motifs and aimed to test this prediction. Classical G4-prediction methods, such as QGRS Mapper or G4Hunter, are based on linear consensus patterns (e.g., (G/C) L) and do not consider deeper and more complex organizational parameters, such as symmetry and periodicity[31], [32], [33], [34]. Trinucleotide symmetry may provide structural benefits for G4 formation, such as optimum loop geometry and optimum register for Hoogsteen base pairing, thereby reducing the energetic barriers to folding. In this study, we sought to identify, categorize, and analyze patterns of trinucleotide symmetry occurrence and locate these elements within the experimentally confirmed G4 motif collection, focusing on motif conservation in the MTOR gene[35]. The MTOR locus is an excellent model, as it contains experimentally confirmed G4 motifs in the first exon that are known to regulate transcription. The choice of this motif is supported by its conservation across primates and other mammals, suggesting its structural and functional relevance[35], [36].

With the help of the quadruplet notation and symmetry matrices of the MTOR gene, we have conducted a comprehensive assessment of the preserved trinucleotide patterns. This study provides an unexplored perspective on trinucleotide symmetry as an indispensable determinant of G4 assembly, further highlighting its value in genomic grammar. Furthermore, we formulated a computational descriptor called the mirror symmetry index (MSI) to quantify the organizational features of G4-forming sequences, namely, the balanced co-occurrence of trinucleotides and their exact reverses within the same strand. This feature can be termed intra-strand mirror symmetry and is distinct from Chargaff’s second parity rule, which measures reverse-complement symmetry across double-stranded DNA.

## Methods

### Data Collection

We retrieved the complete genomic sequences of the MTOR gene from multiple species using the National Center for Biotechnology Information (NCBI) database. Gene sequences from 12 organisms, including primates and commonly used mammalian models, were selected. The chosen species included *Homo sapiens* (humans), *Pan troglodytes* (chimpanzees), *Macaca mulatta* (monkeys), *Gorilla gorilla* (gorillas), *Sus scrofa* (pigs*), Canis lupus familiaris* (dogs), *Mus musculus* (mice), *Rattus norvegicus* (rats), and *Bos taurus* (cattle). We collected and analyzed homologous sequences for the NRAS gene, which served as a comparative reference because of its well-characterized RNA G4 in the 5′ untranslated region (5′ UTR). The downloaded sequences were stored in the FASTA format and processed as needed for downstream analyses. To create a collection of G-quadruplex (G4) motifs that have known properties, we obtained genomic sequences from 14 oncogenes, whose promoter G4s had reached both experimental validation and structural confirmation. The study analyzed G-quadruplex-forming regions which originated from the promoters of c-MYC, VEGF, BCL2, HIF-1α, KRAS, BCR, c-KIT, MET, CARD11, ACC1, HRAS, ATG7, MDM2, and ARID1A. The primary reports and NCBI served as sources for extracting sequences in the top strand, 5′→3′ direction, through the use of coordinates which those studies had established. The reverse-complement strand for each gene was generated to display the complementary sequence in 5′→3′ direction while maintaining the double-stranded DNA structure with its typical antiparallel orientation.

### Identification of Putative Quadruplex-Forming Sequences

QGRS Mapper was used to identify putative quadruplex-forming sequences (PQS) with the canonical search pattern (G/C) □L□□□, (three or more guanines in a row separated by 1-7 nucleotide loops), a pattern that indicates experimentally validated stable G4 motifs. The search was performed with a maximum sequence length of 30 nucleotides for the PQS, a minimum G-score of 20, and allowed loop lengths of 1–7 bases, representing the most significantly enriched PQS type identified in humans. All identified PQS were manually curated to remove overlapping sequences and to confirm the presence of at least three Gs

### Computation of Mirror Symmetry Index (MSI)

The Mirror Symmetry Index (MSI) was computed using Eq. 1:

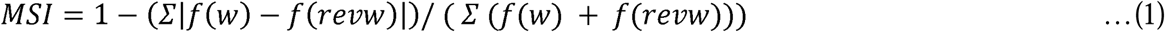

where *f_W_* is the frequency of each distinct trinucleotide, *w* and *f_revW_* is the frequency of the reverse of each trinucleotidew.

To compute the MSI, all overlapping *w* were extracted from the sequences being considered using a sliding window. For a sequence of length *L* this yields *L*-2 nucleotides. For each, *w* its exact reverse *revw* was recorded. The formulation of MSI is analogous to the computation of the S1 symmetry index developed by Huang et al[37]., adapted for using exact reverses instead of reverse complements[3], [38]. Palindromic trinucleotides, such as GGG, GCG, GAG, CGC, and GTG, are self-reversing (*revw* = *w*). While they contribute zero to the numerator of the MSI, they do contribute to the denominator. Their inclusion is consistent with the standard implementation of the S1 symmetry index and was retained throughout the present analysis for methodological consistency and citability. An MSI of 1.0 indicates perfect mirror symmetry, wherein all reverse pairs are perfectly balanced. An MSI of 0 indicates complete asymmetry. Random sequences scored approximately 0.58, and we observed functional G4 sequences with a typical MSI of > 0.80.

## Computation of Non-Palindromic MSI (nMSI)

Non-palindromic MSI (nMSI) was computed using Eq. 2, which excludes palindromic trinucleotides from both the numerator and denominator of the MSI formula:

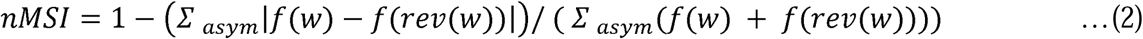

where the summations run exclusively over asymmetric trinucleotide pairs, defined as pairs (w, rev(w)) for which rev(w) ≠ w. Palindromic trinucleotides (those for which the exact reverse equals the trinucleotide itself, such as GGG, CGC, GAG, GTG, and GCG) were excluded from both the numerator and denominator. This exclusion removes a systematic inflation of the denominator caused by palindromic trinucleotides in G-rich sequences, which pushes both wild-type and shuffled MSI values toward 1.0 while contributing no asymmetry information to the numerator. An nMSI of 1.0 indicates perfect mirror symmetry among non-palindromic trinucleotide pairs, and an nMSI of 0 indicates complete asymmetry. If no asymmetric pairs are present (denominator = 0), the nMSI is undefined.

## Results & Discussion

### Highly Conserved PQS in the First Exon of Human MTOR

The primary objective of this study was to perform a comparative analysis of G4 motifs and the underlying trinucleotide symmetry elements across mammalian species. The comparative PQS densities of the MTOR upstream region across nine organisms are shown in Figure S1. Genomic analysis revealed a well-preserved PQS in the first exon of the human MTOR gene across primates[35]. The simultaneous occurrence of the G4 motif in the mRNA transcript (5’ UTR) confirmed that this motif was present in both the DNA and RNA forms, indicating that this motif could potentially be active at the transcriptional and translational levels. The identified sequence (GGGAA GGC GGG CGGT GGGG CAG GGG, 25 bp long) had a G-score of 41 (Table 1). The exonic localization, high G-score, and conservation with DNA and RNA supported the selection of this PQS for further assessment.

**Table 1.**
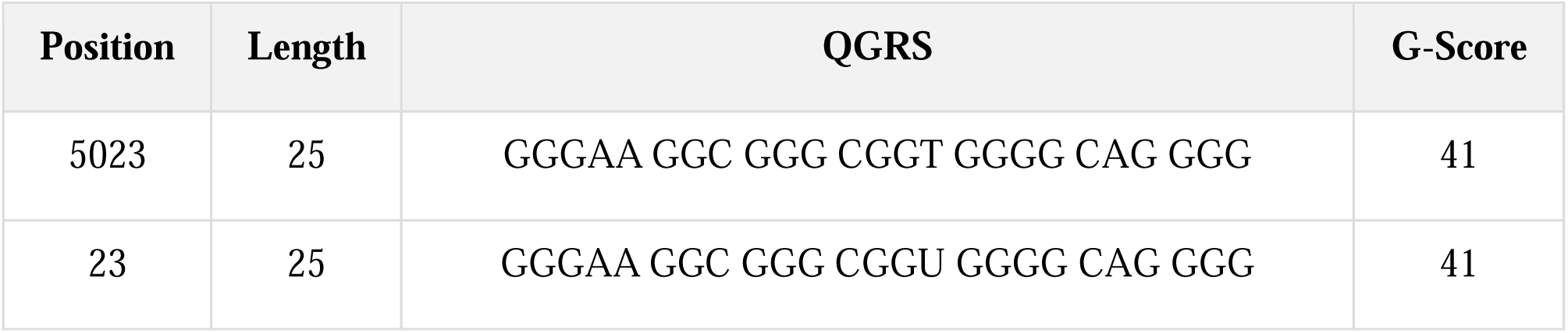
In silico analysis identified PQS motifs present in both the DNA and RNA of the MTOR gene.

### Evolutionary Persistence of G-Tracts and Structural Potential Across Mammalian Species

Multiple sequence alignment of the 12 mammalian species using Clustal Omega showed that the coordinates of 5023 of the human MTOR were shared by all primate sequences analyzed (humans, chimpanzees, gorillas, and macaques) (Figure S2)[39]. The conserved status of the exonic G4 motif among primates implies a healthy selection pressure for preserving the G4-forming potential of the sequence (Figure 1). Although more variation at this locus has been found in non-primate mammals, the overall evidence for G-rich areas and the maintenance of the continuity of G-tracts across mammalian evolution indicate that the ability to form G4 has been conserved throughout evolution[24]. The sequence variants in species did not alter the fundamental structural requirement of sequential guanine repeats with a tolerated loop size. A visualization of trinucleotide symmetry elements in MTOR and NRAS G4-forming sequences is depicted in Figure 2. Furthermore, synonymous and near-synonymous nucleotide changes that do not disrupt the structure have been tolerated. Such conservation of a structural motif despite sequence variation is a molecular hallmark of adaptive selection in G4 biology.

**Figure 1.**
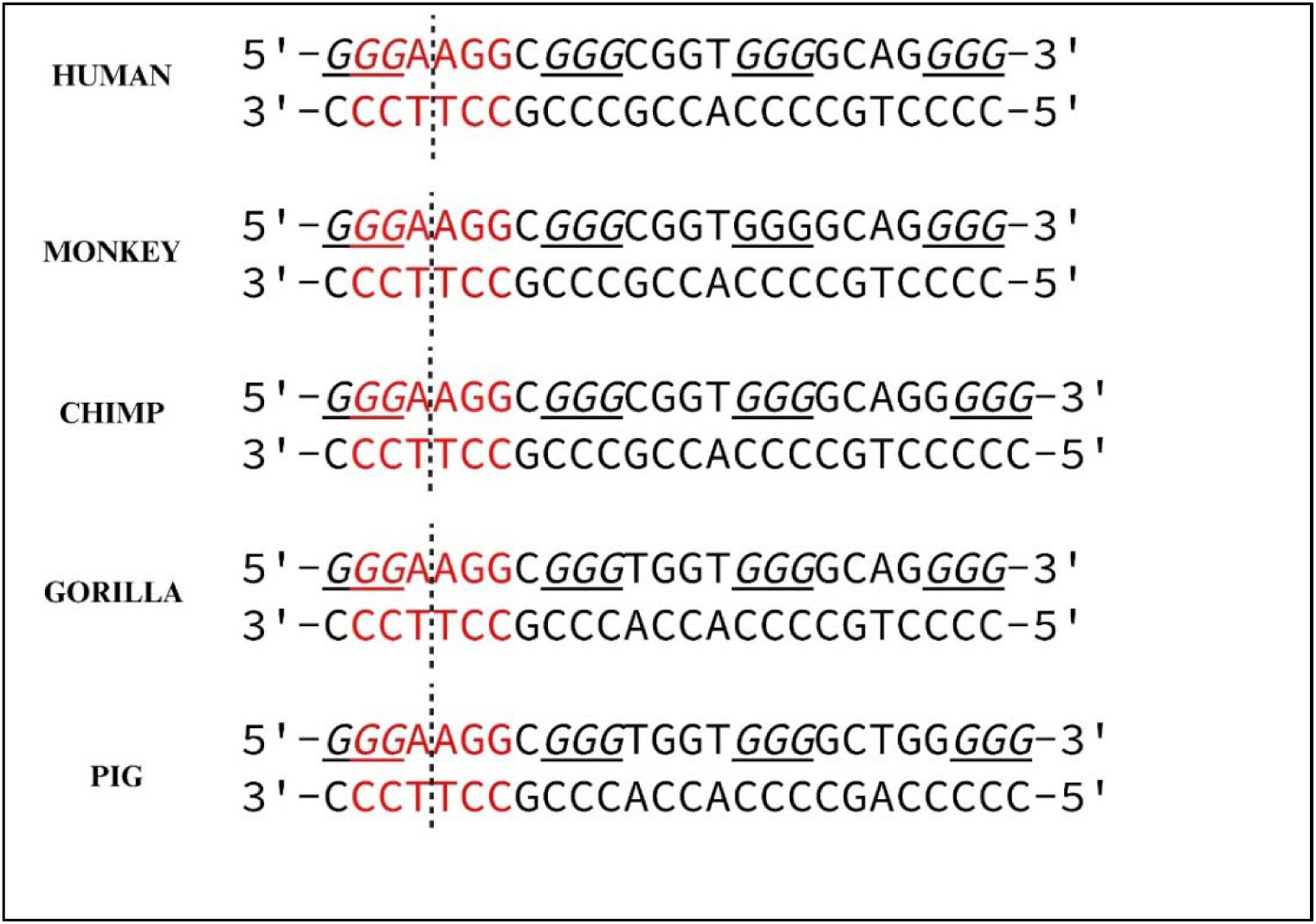
Cross-species symmetry conservation within the MTOR G4 motif.

**Figure 2.**
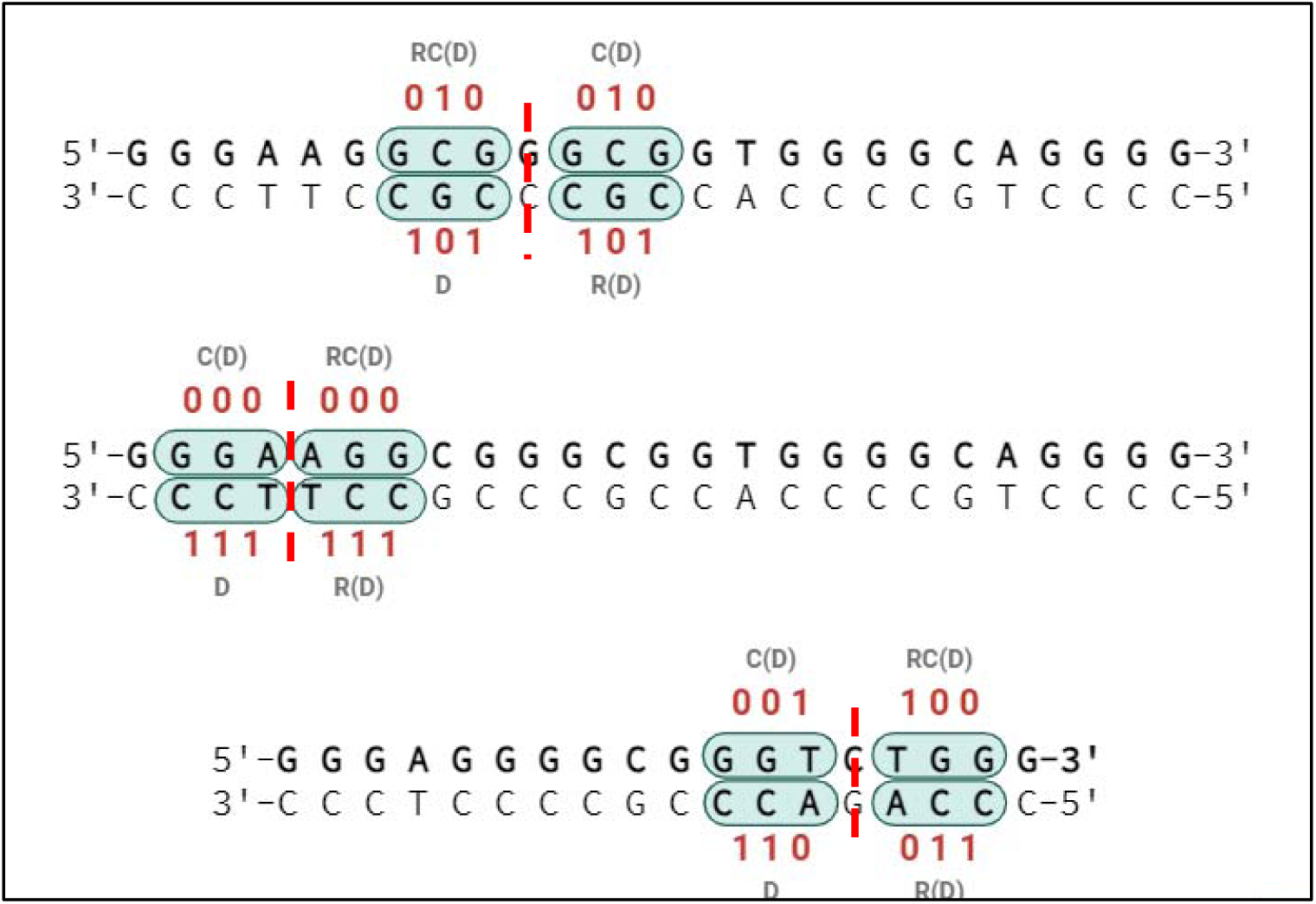
Visualization of trinucleotide symmetry elements within MTOR and NRAS G4-forming sequences.

### Trinucleotide Quadruplet Distribution and Butterfly Symmetry in the MTOR Core

The sequence was next examined by quadruplet matrix classification to determine the trinucleotide composition and symmetry in the conserved MTOR G4 motif and the flanks. The 20 trinucleotide quadruplets were clustered into two major groups: Group I (A+T-rich quadruplets; subgroups Ia, Ib, Ic) and Group II (C+G-rich quadruplets; subgroups IIa, IIb, IIc) (Table S2)[3]. The presence of direct, reverse-complement, complement, and reverse trinucleotides of each quadruplet was examined with specific concern for the mathematical symmetry relationships. Stacked bar histograms of the quadruplet frequency distributions of primates species provided several important results: (1) There are always mirror symmetry patterns in both A+T-rich and C+G-rich matrices of all primates species studied; (2) Relative frequencies of direct trinucleotides were very similar in all matrices groups, even though there was slight absolute nucleotide differences; (3) Butterfly symmetry exists, showing that the frequencies of reverse complementary trinucleotides are very similar.

Interestingly, the levels of symmetry of the MTOR G4 motif and adjacent regions were higher than those of the background exonic sequences of the same gene[4], [40]. The amplification of mirror-symmetric trinucleotides (especially in the G-rich core of the motif) contrasts with the overall asymmetry of codon-biased exons, indicating that functional G4 sequences are exposed to the selection of symmetry patterns beyond the demands of protein-coding.

### Symmetry as a Universal Determinant of Biologically Relevant Oncogenic G4s

To evaluate whether trinucleotide symmetry is a common property of biologically relevant G4s, we evaluated the experimentally validated G4-binding sequences of the oncogenes c-MYC, VEGF, BCL2, and KRAS[41], [42], [43], [44]. The selection of these genes was based on their experimentally studied G4 motifs, which play a role in transcription modulation, serve as therapeutic targets, and are present in a wide variety of genomic contexts (promoter vs. UTR localization). As presented in Table 2, the sequences were grouped by the reported G4 topology (parallel, anti parallel, mixed). For each gene, the top strand (5′→3′) and its reverse complement (bottom strand, also shown as 5′→3′) are presented. Red, green, and pink highlights mark trinucleotide elements that participate in mirror symmetry relationships (e.g., direct vs. reverse complement or purine pyrimidine mirror repeats). Different colors are used to distinguish distinct symmetry elements within the same G4 sequence, with each color representing mirror-symmetry contributions. The canonical c-MYC promoter G4 was analyzed, and a remarkable reflection pattern on trinucleotide composition was observed (Table 2). Purine-pyrimidine mirror repeats (GTG (direct) and CAC (reverse-complement)) and GAG (direct) and CTC (reverse-complement) trinucleotides were overrepresented in the sequence. These trinucleotides are symmetry subgroups IIa and Ic, respectively, and the frequencies of these trinucleotides are almost mirror symmetric. The recurring nature of the guanine-rich trinucleotides is interrupted by symmetric purine-pyrimidine dyads, forming a structure whose trinucleotide structure is optimized to form a stable G-tetrad by maximizing the loop geometry and electrostatic interactions. The BCL2 G4 motif also exhibits high trinucleotide symmetry, with direct and reverse complement symmetries. It was found to be enriched in symmetric trinucleotides (Group Ic and IIc subgroups, which have self-complementary sequences) and indicated that the architecture of the sequence is enforced to maintain a symmetric structure between the strands. VEGF and KRAS promoter G4s have mirror-symmetric trinucleotide patterns with slight differences in the extent of symmetry and enrichment of particular trinucleotides. VEGF is specifically enriched in symmetrical trinucleotides (Ic and IIc subgroups), whereas KRAS has a more complicated structure with several interleaved symmetry elements.

**Table 2.**
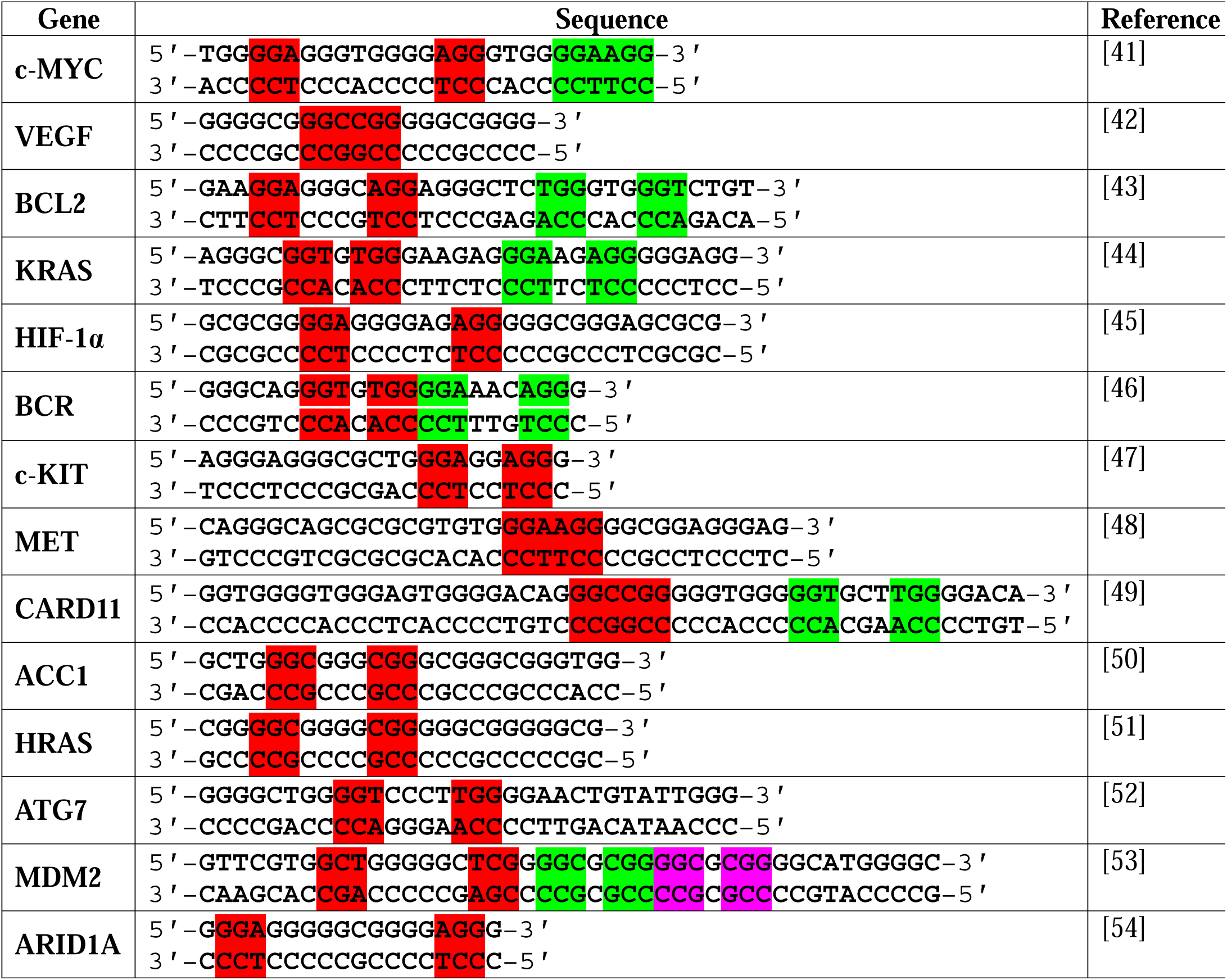
Prevalent symmetries within various biologically relevant oncogenic G4s.

### Interspecies Comparison of Trinucleotide Frequencies of MTOR

Next, we examined the trinucleotide frequencies across the entire MTOR locus. Interestingly, asymmetries in exonic trinucleotide frequencies are compensated for by alterations in non- coding sequences at the ends[4], [11]. In particular, the increased frequency of A+T-rich quadruplets in exons was accompanied by an inverse decrease in C+G-rich quadruplets (Figure S3). This inverse characteristic helps maintain the overall number of nucleotides in the chromosome to a constant while facilitating the asymmetries caused by codon bias, which are required to allow protein synthesis in exons.

Notably, the extent of these asymmetries varies between species, depending on their codon usage preferences, reflecting species-specific adaptation to translational efficiency and tRNA availability[40]. In contrast, the intronic and intergenic regions flanking the MTOR gene maintain higher degrees of trinucleotide symmetry, consistent with the preservation of Chargaff’s second parity rule (CSPR) in non-coding DNA[4]. The inverse proportional relationship between A+T-rich and C+G-rich trinucleotide quadruplets, observed to be preserved in the overall MTOR locus despite local coding region asymmetries, indicates that compensatory nucleotide distributions in non-coding regions maintain genome-wide symmetry while permitting a functional coding region bias (Figure 3). This framework explains why orthologous MTOR genes across species show both conservation of certain symmetrical trinucleotide patterns (reflecting shared functional constraints) and species-specific deviations (reflecting divergent codon usage optimization), with the magnitude of deviation being inversely correlated with the proportion of non-coding sequence length flanking the gene. This conservation pattern extends from mammals to other vertebrates and even to more distantly related species, indicating a strong selective pressure to maintain these structural elements.

**Figure 3.**
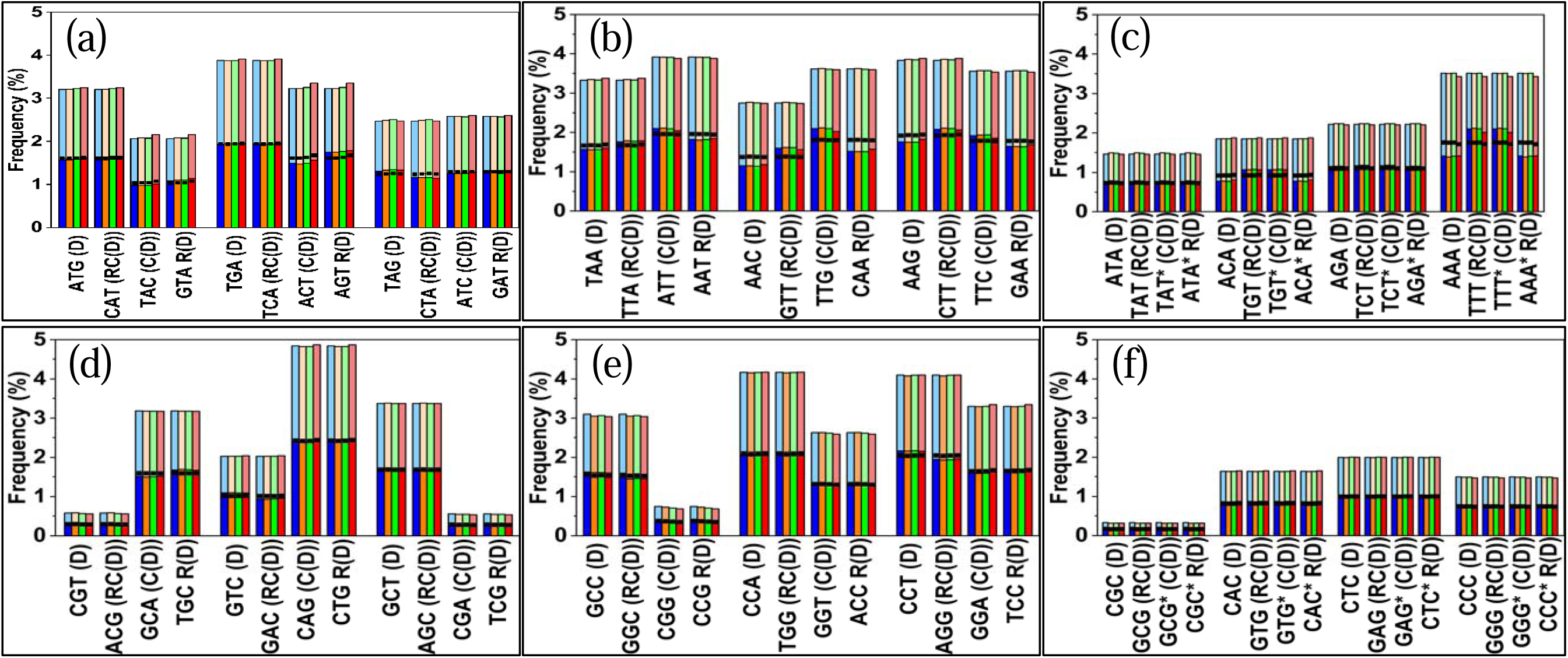
Comparison of quadruplet matrices for the mTOR gene in four primate species. Stacked bar histograms showing the frequency (%) of trinucleotides belonging to symmetry groups (a) Ia, (b) Ib, (c) Ic (A+T-rich matrices), and (d) IIa, (e) IIb, (f) IIc (C+G-rich matrices) in the mTOR locus of *Homo sapiens* (human, blue), *Gorilla gorilla* (gorilla, orange), *Pan troglodytes* (chimpanzee, green), and *Macaca mulatta* (monkey, red). For each species, frequencies are displayed for both the top strand (5′→3′, darker shade) and bottom strand (reverse-complement, lighter shade). The black horizontal line within each bar represents the mean frequency across the four species for that trinucleotide in the corresponding strand. In each quadruplet, the frequencies of direct (D) and reverse complement (RC) trinucleotides are equal (D = RC), and the frequencies of complement (C) and reverse (R) trinucleotides are equal (C = R), consistent with butterfly symmetry.

Notably, the region of the MTOR G4 motif has a slightly different pattern: there is a certain asymmetry of the exon with the presence of G-tract and the usage of codons, but the flanking sequences and the G4 motif itself are less asymmetrical than non-functional exonic areas in the same gene. This implies that functional G4 sites face two selective forces: (1) the ability to retain protein-coding ability by using codons and (2) the ability to retain trinucleotide symmetry to allow G4 formation and regulation. The outcome is a series in which symmetry has been locally raised with respect to the neutral exonic background, indicating that this higher-order organizational property has been selected.

### Selective Pressure for Localized Symmetry within Asymmetric Protein-Coding Sequences

The molecular basis for the violation of CSPR in coding DNA is justified based on codon usage bias[40]. The genetic code is degenerate, and several codons encode an amino acid. Some codons are more favored by different organisms than the alternatives that are synonymous, which have been motivated by the presence of tRNAs, efficacies in translation, and other translational optimization processes. A strong overrepresentation of A-codons, especially AAA and AAG (codons that encode lysine) and AAC (codons that encode asparagine), is widespread in most organisms, including humans. This bias in favor of adenine produces an imbalance in the frequencies of the nucleotides in the coding sequences: the strand of coding sequences contains more A residues than T, and its complementary strand contains more T residues than A.

In contrast, at the genome-wide or chromosomal level, CSPR is largely maintained by a compensatory process known as inverse proportionality[4], [13]. Variations in A+T-rich quadruplet counts (Figure 3a, 3b, and 3c) in the coding regions are counteracted by counter-variations in C+G-rich quadruplet counts (Figure 3d, 3e, and 3f) in the coding sequences or adjacent regions. This trade-off is an evolutionary constraint that preserves functional integrity and genomic stability over time. The MTOR G4 motif and other functional G4s pose a riddle in this context: they are found in coding regions (exons) but still have high trinucleotide-symmetry levels despite the asymmetries of codons on which the codon bias theory relies. We suggest that this local asymmetry may indicate a process of selective pressure to maximize the G4 structure. Specifically, mirror and reverse-complement arrangements of trinucleotide composition symmetries are expected to be structurally beneficial for forming G-quadruplexes. These symmetric trinucleotide architectures can minimize the energetic penalty for quadruplex assembly by partially pre-organizing the nucleic acid strand into conformations that are already predisposed to G4 formation, thereby reducing the entropic cost associated with G4 assembly. Furthermore, purine–pyrimidine mirror symmetries might influence a favorable electrostatic environment for Hoogsteen hydrogen bonding and the orthogonal geometric orientation of loop regions connecting neighboring G-tetrads. These symmetries may also enhance the optimal coordination of monovalent cations in the central G-quadruplex channel, which is an important factor in structural stabilization. Finally, as trinucleotide sequences are known to have certain biased loop conformations, the symmetric repetition of such sequences can further stabilize the structure through stabilized loop–loop and loop–core interactions, increasing the stability and robustness of the G-quadruplex.

### MSI of G4-Motifs in Oncogene Promoters

Prominent G4 prediction tools, such as QGRS Mapper and G4Hunter, use either the pattern-matching algorithm or the empirical scoring algorithm, which inherently excludes the higher-order organizational context[31], [32]. These tools are designed to search for linear consensus sequences without adding information regarding trinucleotide distribution and symmetry. Therefore, they yield large false-positive and false-negative rates, predicting non-functional G-rich sequences as G4s and missing non-canonical and biologically relevant G4s, respectively. Trinucleotide symmetry analysis should be part of future computational approaches as a prioritization layer.

To quantitatively develop trinucleotide symmetry metrics that can be correlated with G4 formation, we calculated the MSI of a panel of 14 experimentally characterized G4 motifs from oncogene promoters. MSI calculations were performed as described in the Methods section, and the results are shown in Table 3. The length (L) of the promoter sequences studied varied from 18 (ARID1A) to 49 (CARD11), spanning short parallel promoter G4s and longer loop-rich motifs.

**Table 3.**
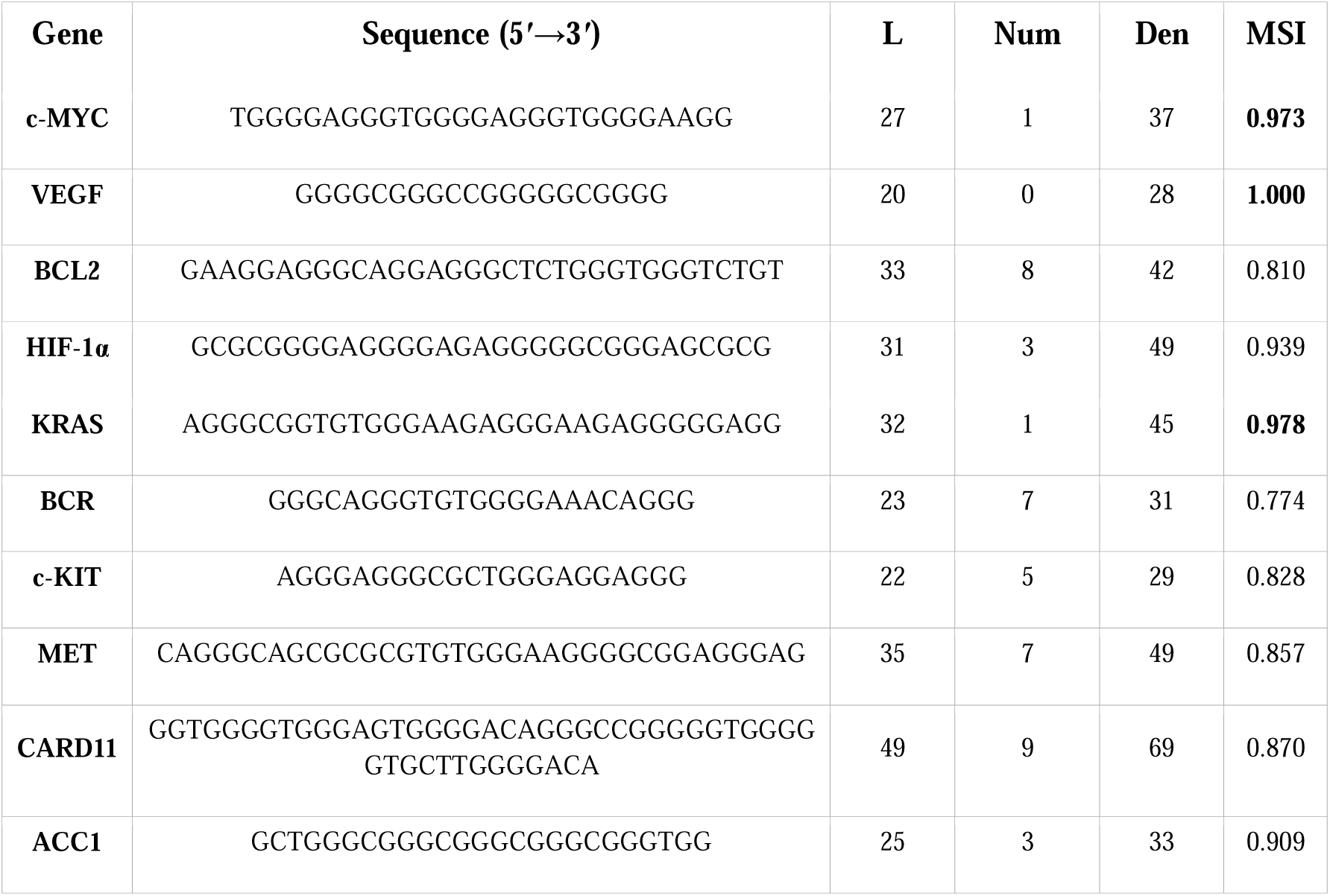
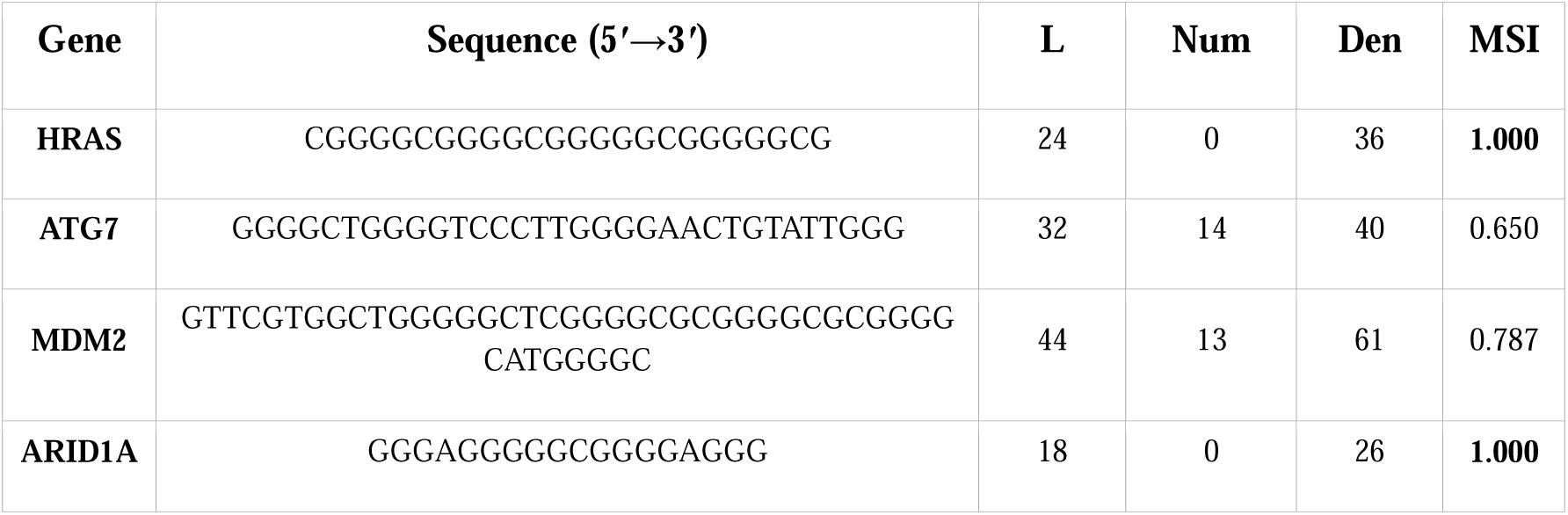
MSI values of G4-motifs in oncogene promoters.

Notably, the MSI of most oncogene promoter G4s exceeded 0.80 (mean 0.884), and the highest values (MSI ≥ 0.950, shown in bold) were obtained for c-MYC (0.973), KRAS (0.978), VEGF (1.000), HRAS (1.000), and ARID1A (1.000). A minority of motifs fell below the 0.80 mark, namely ATG7 (0.650), BCR (0.774), and MDM2 (0.787); these are the longest and most loop-rich sequences in the panel, in which extended asymmetric loop regions dilute the balanced core symmetry. The high MSI of G-rich sequences raises the question of exceeding chance expectations. Accordingly, we subjected each G4-forming sequence to 1000 mononucleotide-composition-preserving random shuffles (uShuffle, k=1)[55]. The Fisher-Yates permutation preserves exact nucleotide counts while randomizing the order, generating a null distribution of MSI values for each sequence. All 15,000 shuffled sequences were verified to preserve the mononucleotide composition. We did not use trinucleotide-preserving shuffles because MSI is computed entirely from trinucleotide frequencies. A trinucleotide shuffle that preserves trinucleotide counts will, by mathematical necessity, produce the same MSI as wild-type sequences. This is merely a mathematical identity, not a biological result. Thus, mononucleotide-preserving shuffle analysis is the only informative null model for MSI. A comparison of the wild-type MSI and mononucleotide-preserving shuffled MSI is shown in Table 4.

Tables□4 and 5 depict two complementary analyses of the 14 WT G4-forming sequences against their mononucleotide-composition-preserving shuffle null distributions. Table□4 reports the MSI (Eq.□1) while Table□5 reports the non-palindromic MSI (nMSI, Eq.□2). Non-palindromic MSI excludes palindromic trinucleotides whose exact reverse equals themselves, such as GGG, GCG, GAG, CGC, and GTG, from both the numerator and denominator. This exclusion addresses a systematic confound in these G-rich sequences, where palindromic trinucleotides account for a large fraction of the MSI denominator while contributing zero to the numerator. This pushes both the wild-type and shuffled MSI toward the same ceiling of ≈1.0, preventing the wild-type from distinguishing itself from its compositional background. The nMSI removes this shared ceiling and focuses the index on asymmetric trinucleotide pairs that carry genuine structural information about the DNA molecule. In the MSI analysis shown in Table□4, the KRAS promoter G4 is the only sequence that reaches individual significance against its null distribution (Δ□=□+0.106, p□=□0.042), and the same sequence attains the strongest nMSI signal in Table□5 (Z□=□+1.906, p□=□0.0349). For the remaining sequences, no individual test reached p□≤□0.05, which reflects a fundamental statistical power limitation: the motifs yield relatively few trinucleotide observations, and the constrained mononucleotide-preserving shuffle space for such G-rich sequences produces narrow null distributions. Two sequences, ATG7 and MDM2, scored below their shuffle means on both metrics (Table□4: Δ□=□−0.085 and −0.075; Table□5: Z□=□−1.202 and −1.557), owing to their inherently asymmetric loop trinucleotide compositions. To assess the collective significance of the trend, we applied Stouffer’s combined Z-score method, pooling the 14 individual nMSI Z-scores: Z□_combined_□=□1.197, p□=□0.116 (n□=□14). The corresponding trend for the original MSI is comparably weak, indicating that neither index reaches collective significance under this conservative per-sequence null model. Overall, eight of the 14 sequences showed nMSI(WT)□>□nMSI shuffle mean, and three sequences (VEGF, HRAS, and ARID1A) achieved the maximum nMSI of 1.000, indicating perfect mirror symmetry in their non-palindromic trinucleotide pairs. The ATG7 and MDM2 sequences were the clearest outliers (nMSI□=□0.300, Z□=□−1.202; and nMSI□=□0.435, Z□=□−1.557, respectively): the specific loop architectures of these long, loop-rich motifs generated non-palindromic trinucleotide pairs that were highly imbalanced, yielding genuinely lower non-palindromic mirror symmetry. This is a biologically interpretable result reflecting the structural complexity of these G4s, not a contradiction of the mirror symmetry hypothesis. HIF-1α is another instructive case: although it retains a high MSI (0.939) and a moderate nMSI (0.667), its experimentally confirmed G4-abolished mutant coincidentally generates a more palindrome-rich sequence, so that the mutant outscores the wild-type on both indices (Table□6). This reversal reflects a known limitation of scrambled-sequence mutant controls, rather than a failure of the metric.

**Table 4.** Wild-type MSI compared to the null distribution from 1000 mononucleotide-preserving shuffles.

| Gene | WT MSI | Mean (shuffled) | SD | Min | Max | $\Delta$ (WT-Shuf) | p-value | Verdict |
| --- | --- | --- | --- | --- | --- | --- | --- | --- |
| c-MYC | **0.973** | 0.911 | 0.055 | 0.703 | 1.000 | **+0.062** | 0.213 | **Trend WT > rand** |
| VEGF | 1.000 | 0.981 | 0.019 | 0.933 | 1.000 | +0.019 | 0.463 | WT ≈ rand |
| BCL2 | 0.810 | 0.751 | 0.080 | 0.474 | 0.978 | **+0.059** | 0.256 | **Trend WT > rand** |
| HIF-1α | 0.939 | 0.903 | 0.057 | 0.619 | 1.000 | +0.036 | 0.247 | WT ≈ rand |
| KRAS | 0.978 | 0.872 | 0.055 | 0.634 | 1.000 | **+0.106** | **0.042** | **Trend WT > rand** |
| BCR | 0.774 | 0.775 | 0.088 | 0.519 | 1.000 | -0.001 | 0.545 | WT ≈ rand |
| c-KIT | 0.828 | 0.833 | 0.074 | 0.593 | 1.000 | -0.005 | 0.580 | WT ≈ rand |
| MET | 0.857 | 0.807 | 0.070 | 0.512 | 1.000 | +0.050 | 0.263 | WT ≈ rand |
| CARD11 | 0.870 | 0.850 | 0.052 | 0.625 | 0.973 | +0.020 | 0.391 | WT ≈ rand |
| ACC1 | 0.909 | 0.906 | 0.058 | 0.722 | 1.000 | +0.003 | 0.539 | WT ≈ rand |
| HRAS | 1.000 | 0.985 | 0.016 | 0.944 | 1.000 | +0.015 | 0.491 | WT ≈ rand |
| ATG7 | 0.650 | 0.735 | 0.087 | 0.389 | 0.977 | **-0.085** | 0.842 | **WT < rand**<br>□ |
| MDM2 | 0.787 | 0.862 | 0.052 | 0.655 | 0.984 | **-0.075** | 0.916 | **WT < rand**<br>□ |
| ARID1A | 1.000 | 0.937 | 0.055 | 0.815 | 1.000 | **+0.063** | 0.256 | **Trend WT > rand** |

**Table 5.**
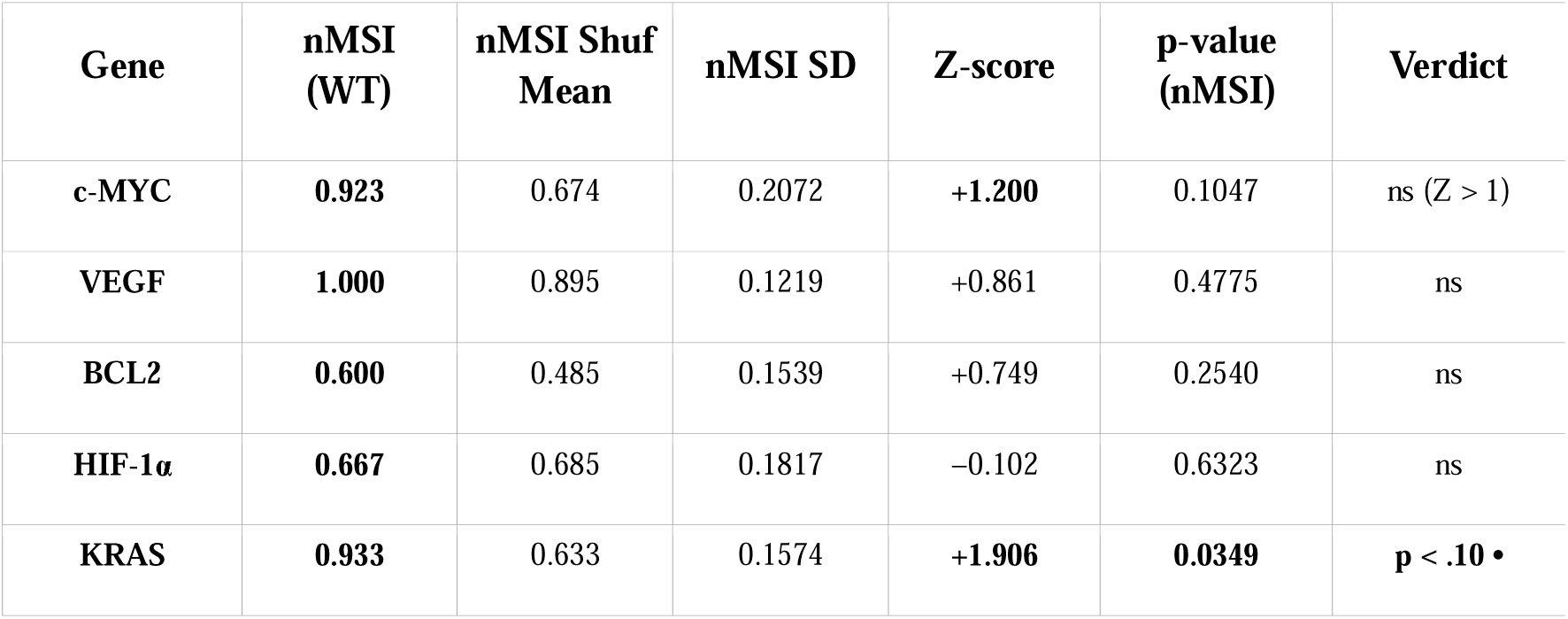
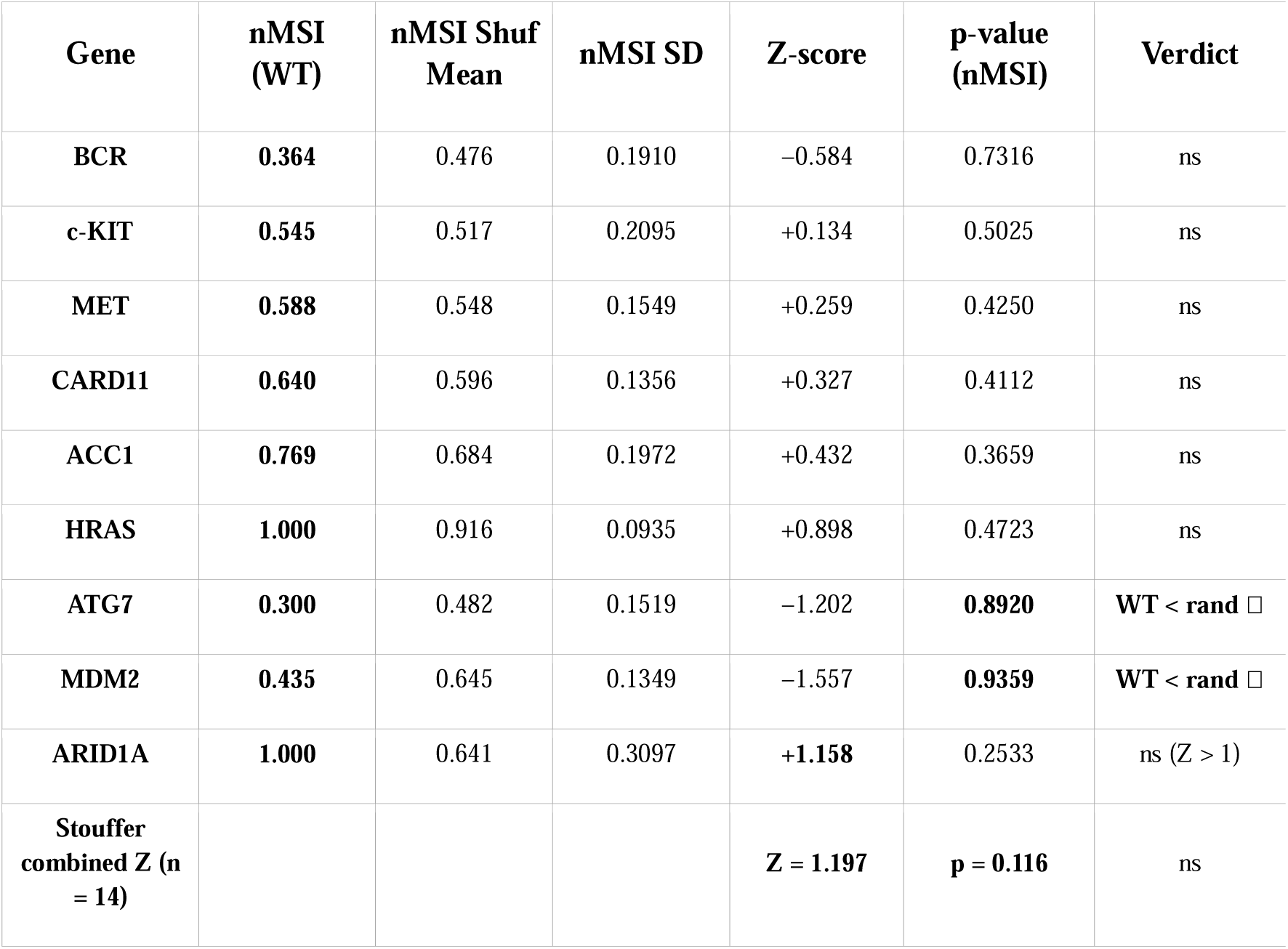
Non-palindromic MSI (nMSI) compared to the null distribution from 10,000 mononucleotide-preserving shuffles per sequence.

| Gene | nMSI (WT) | nMSI Shuf Mean | nMSI SD | Z-score | p-value (nMSI) | Verdict |
| --- | --- | --- | --- | --- | --- | --- |
| c-MYC | 0.923 | 0.674 | 0.2072 | **+1.200** | 0.1047 | ns (Z > 1) |
| VEGF | 1.000 | 0.895 | 0.1219 | +0.861 | 0.4775 | ns |
| BCL2 | 0.600 | 0.485 | 0.1539 | +0.749 | 0.2540 | ns |
| HIF-1α | 0.667 | 0.685 | 0.1817 | -0.102 | 0.6323 | ns |
| KRAS | 0.933 | 0.633 | 0.1574 | **+1.906** | **0.0349** | **p < .10 •** |
| BCR | 0.364 | 0.476 | 0.1910 | -0.584 | 0.7316 | ns |
| c-KIT | 0.545 | 0.517 | 0.2095 | +0.134 | 0.5025 | ns |
| MET | 0.588 | 0.548 | 0.1549 | +0.259 | 0.4250 | ns |
| CARD11 | 0.640 | 0.596 | 0.1356 | +0.327 | 0.4112 | ns |
| ACC1 | 0.769 | 0.684 | 0.1972 | +0.432 | 0.3659 | ns |
| HRAS | 1.000 | 0.916 | 0.0935 | +0.898 | 0.4723 | ns |
| ATG7 | 0.300 | 0.482 | 0.1519 | -1.202 | 0.8920 | WT < rand □ |
| MDM2 | 0.435 | 0.645 | 0.1349 | -1.557 | 0.9359 | WT < rand □ |
| ARID1A | 1.000 | 0.641 | 0.3097 | +1.158 | 0.2533 | ns (Z > 1) |
| Stouffer combined Z (n = 14) |  |  |  | Z = 1.197 | p = 0.116 | ns |

The comparison of the MSI of each wild-type G4-forming sequence against its experimentally validated G4-abolished mutant counterpart would constitute a more biologically meaningful validation than shuffling. Mutant sequences carrying substitutions (G→T or G→C at key G-run positions) confirmed by circular dichroism (CD) spectroscopy, NMR, or cellular G4-ChIP assays to abolish G4 formation were drawn wherever possible from the primary literature (Table 6). This comparison tests whether the MSI is specifically sensitive to the sequence features required for G4 folding; if the MSI reflects G4-forming capacity, wild-type sequences should consistently score higher than their G4-negative mutants. A comparison of the MSI of wild type and G4-abolished mutants is presented in Table 6. WT MSI exceeded mutant MSI in 12 of 14 paired comparisons (mean ΔMSI = +0.089), and the effect was even larger on the non-palindromic index (WT nMSI > mutant nMSI; mean ΔnMSI = +0.192). A one-tailed sign test on the 12/14 MSI outcome gave p = 0.0065, confirming a robust directional association between mirror symmetry and G4-forming capacity. The strongest MSI effects were observed for VEGF (Δ = +0.364), c-MYC (+0.246), and HRAS (+0.176). Two genes, BCL2 (Δ = −0.123) and HIF-1α (Δ = −0.061), showed a reversed pattern (WT MSI < mutant MSI); in both cases, the scrambled or substituted mutant coincidentally introduced a more balanced trinucleotide composition, a recognized limitation of scrambled-sequence controls rather than a biological failure of the index.

**Table 6.**
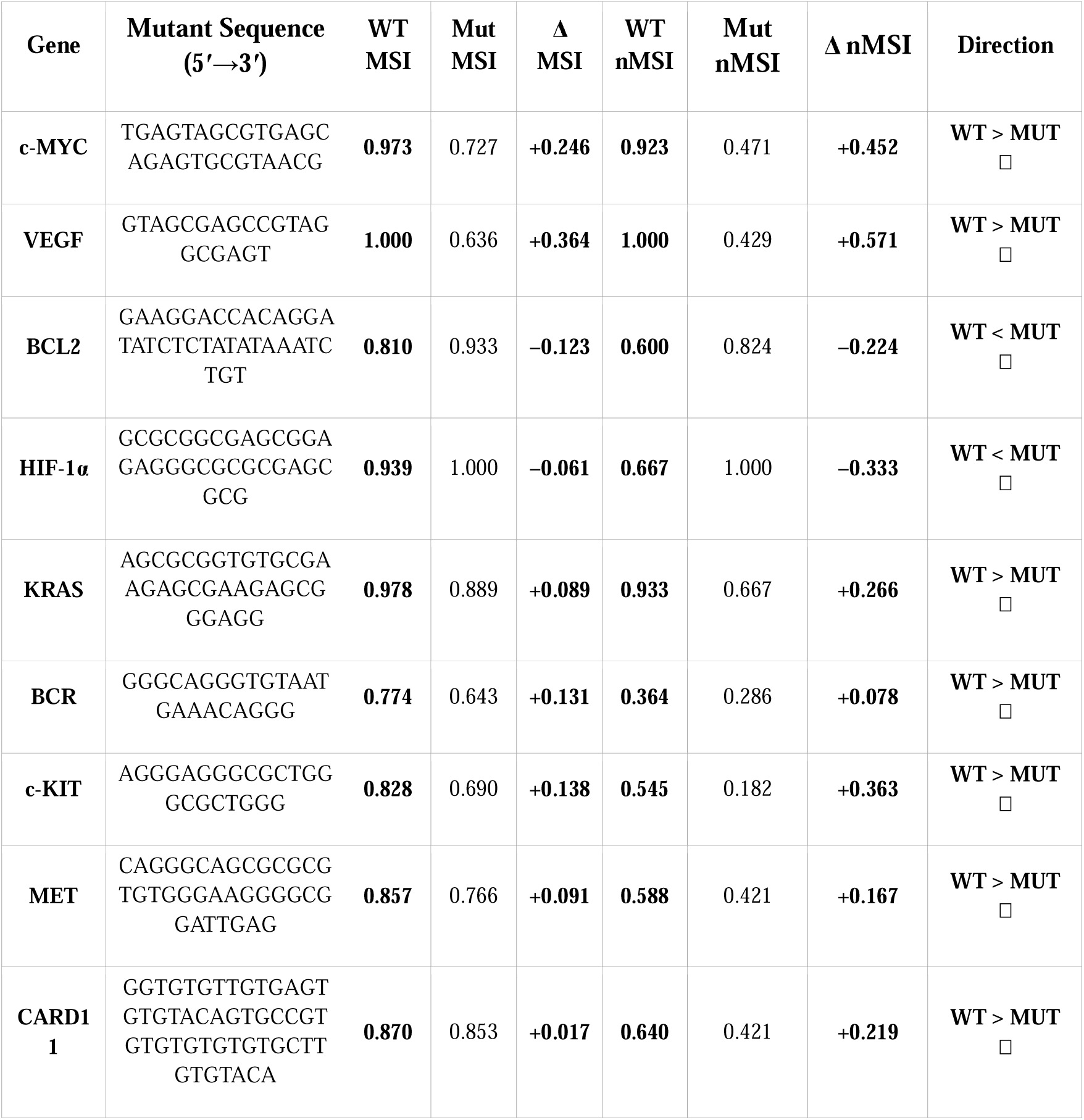
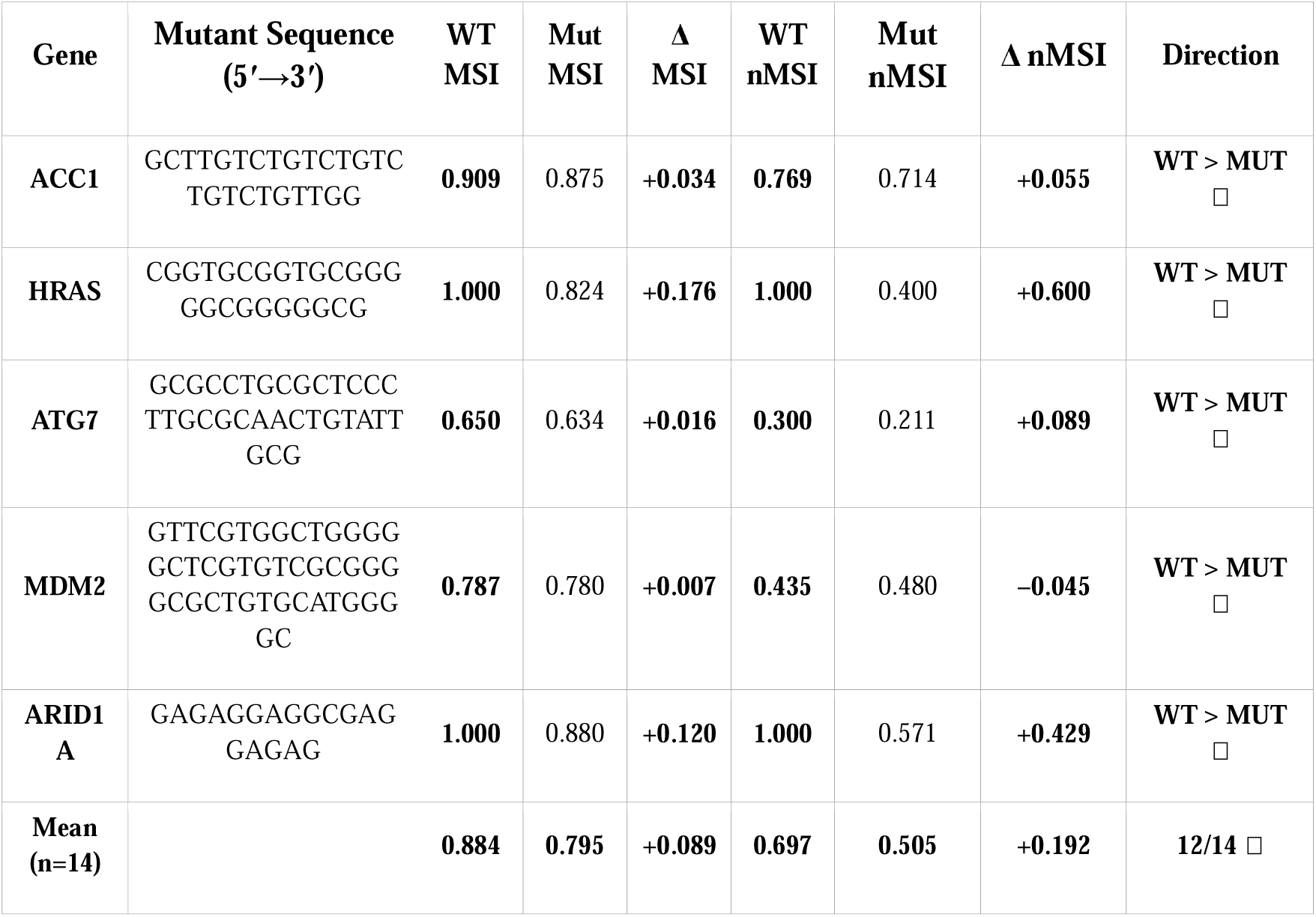
MSI and nMSI comparison of wild-type vs. G4-abolished mutant.

Next, we tested MSI on recently published G4 sequences to compare the MSI of the reported functional G4s against PQS’ in the same gene and against experimental negative controls. In this regard, we chose G4 sequences that have recently been reported in the B-MYB promote[56]r. Miranda et al. (2024) identified six G4-forming sequences in the B-MYB promoter (Chromosome 20), spanning a range of G4Hunter (G4H) scores from 1.12 to 2.94. Sequences with high G4H scores (B-MYB 25RC and B-MYB 22R, scores 2.94 and 2.91, respectively) represent the strongest G4-forming candidates, whereas those with low G4H scores (1.12–1.20) represent weak candidates. A comparison of the MSI scores for these PQS is presented in Table 7.

**Table 7.**
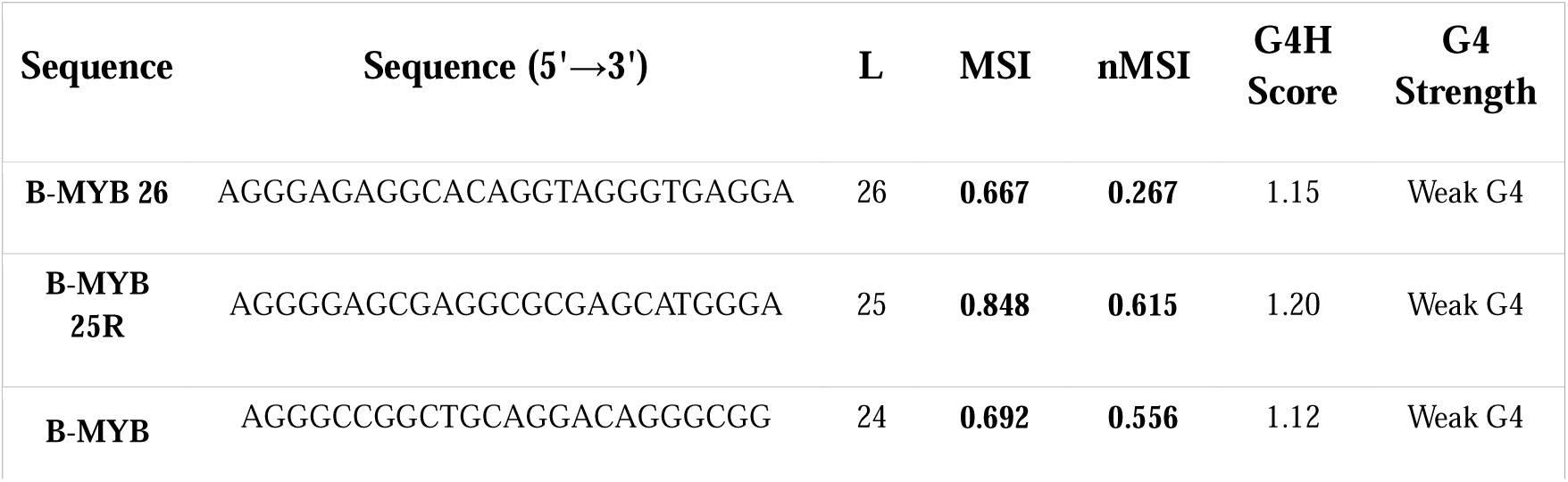
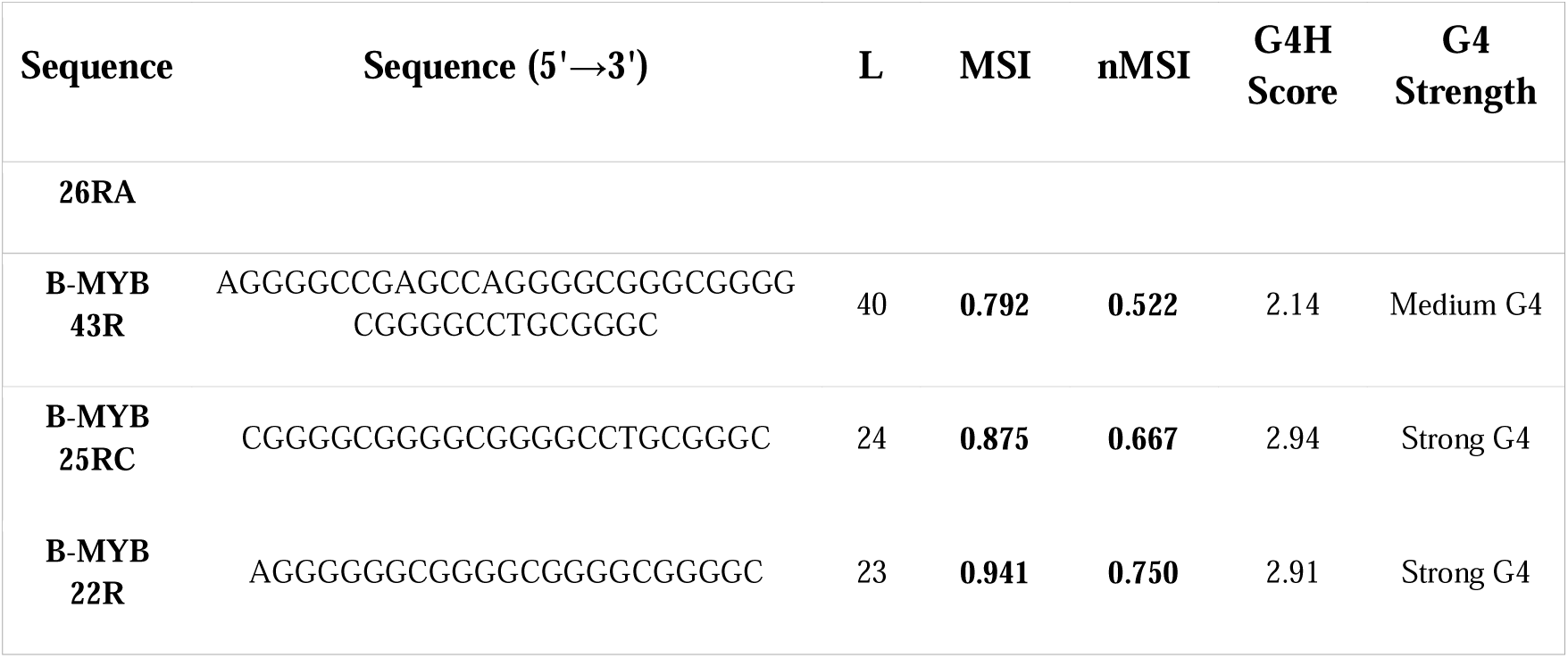
MSI and nMSI of PQS in the B-MYB promoter.

As shown in Table 7, High G4H sequences scored higher MSI (mean = 0.908) than low G4H sequences (mean = 0.736). MSI positively correlated with G4 propensity across this series, providing evidence that mirror symmetry is associated with G4-forming potential within a single gene. Esain-Garcia et al. used CRISPR genome editing to abrogate endogenous G4 structure formation at the MYC locus[57]. Wild-type MYC and KRAS swap sequences to form G4 structures, as verified by CD spectroscopy. Multiple mutant variants (MYC MUT, MUT CORE, and MUT MIN) were confirmed by CD to lack G4 formation, while MYC FLIP placed the G4 on the template strand. We tested the MSI on these wild-type and mutant sequences, and the results are presented in Table 8.

**Table 8.**
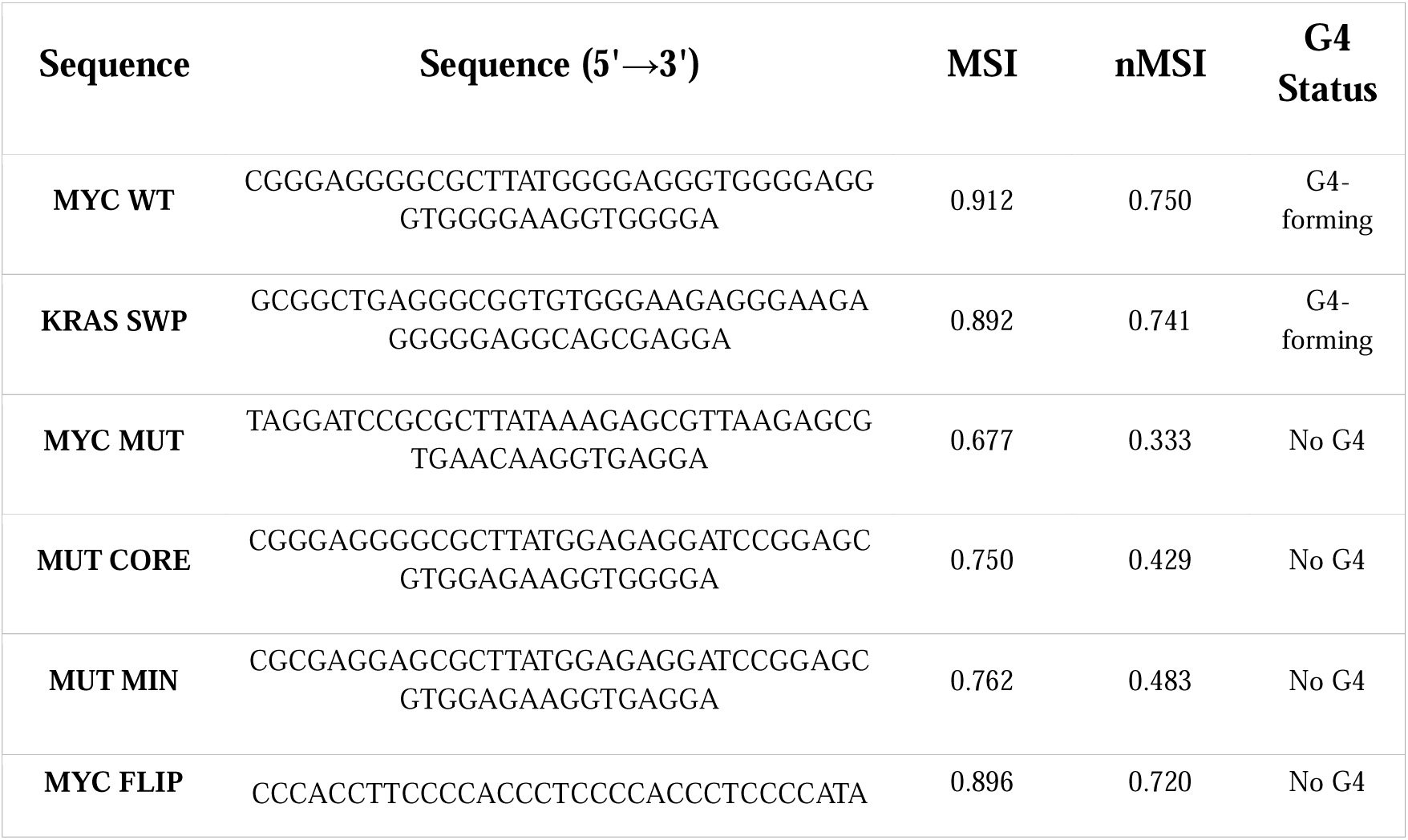
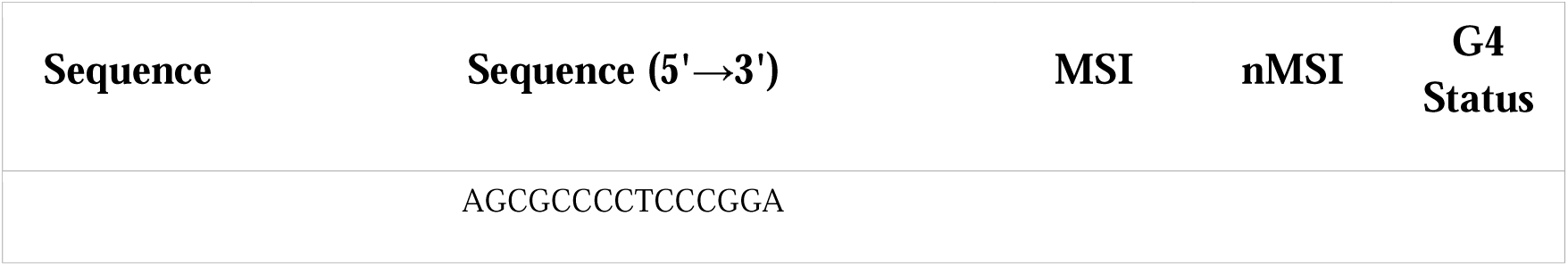
MSI and nMSI of MYC CRISPR-edited sequences.

As shown in Table 8, G4-forming sequences (MYC WT and KRAS SWP) had a mean MSI of 0.902, whereas non-G4 sequences (MYC MUT, MUT CORE, and MUT MIN) had a mean MSI of 0.730. Δ = +0.172. The lengths of all these sequences (48 nt) were the same. MYC FLIP (0.896) was an exception in terms of MSI. It retains a high MSI because its trinucleotide composition is identical to that of MYC WT (only the strand is switched), illustrating that MSI measures sequence composition, not strand orientation. In another recent study, Chen et al. used G4-ChIP-qPCR to confirm G4 folding in living human cells using GPX4 and CLIC4 promoter sequences[58]. G4-negative mutants were constructed by replacing key G bases with lowercase/mutant residues, and the loss of the G4 signal was confirmed by G4-ChIP.

As shown in Table 9, G4-forming sequences scored higher MSI (mean = 0.823) than G4-negative mutants (mean = 0.732) with Δ = +0.091. While the modest absolute differences reflect the shorter sequence lengths, the directional result is consistent across both gene pairs and with our hypothesis. Based on these assessments, we demonstrated the ability of MSI to consistently differentiate G4-forming sequences from non-G4 controls across multiple independent datasets and validation strategies. The comparison of G4-abolished mutants of identical length (Tables 6, 7, and 8, where sequences are length-matched) provides a length-controlled validation of MSI. The cross-validation of MSI across three independent external datasets, together with the wild-type versus G4-abolished mutant sign test over 14 paired comparisons, provides a global assessment that is robust to multiple hypothesis-testing concerns. Aside from the KRAS promoter G4 (p = 0.042 on MSI; p = 0.0349 on nMSI), individual shuffle-based p-values did not reach significance, reflecting a power limitation imposed by the short sequence lengths and extreme G-richness of these motifs, rather than the absence of a biological signal. The wild-type versus G4-abolished mutant comparison, however, yielded p = 0.0065 by a one-tailed sign test on 12 of 14 concordant pairs — a single global test not subject to multiple comparison correction — confirming a robust directional association between mirror symmetry and G4-forming capacity.

**Table 9.**
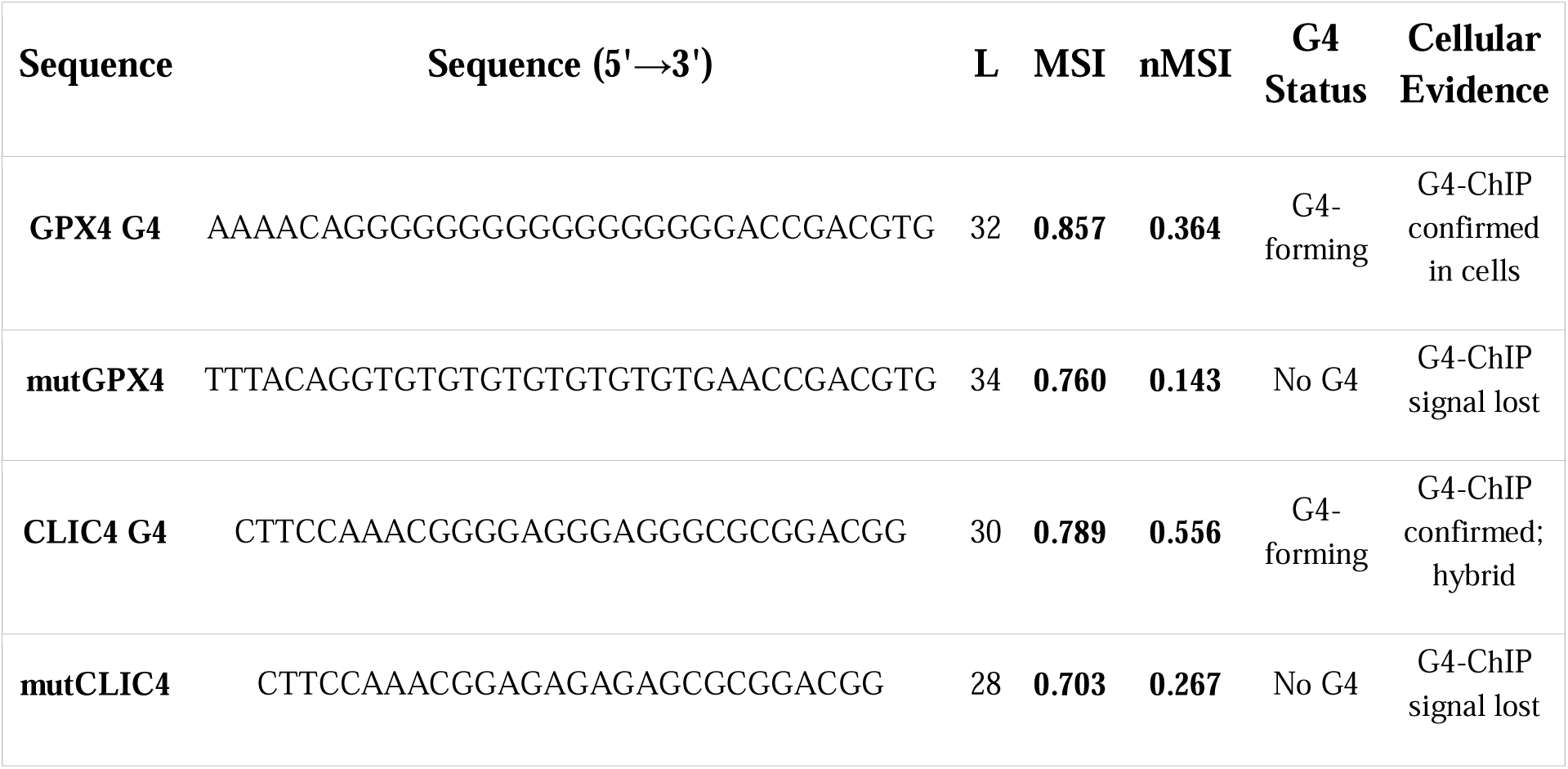
MSI of GPX4 and CLIC4 G4-forming sequences and their G4-negative mutants.

### nMSI ≥ 0.50 Correlates with G4-Folding Propensity

Although the per-sequence shuffle tests provided only a non-significant collective trend for nMSI (Stouffer Z = 1.197, p = 0.116), the wild-type versus mutant comparison established a clear directional signal. Therefore, we examined whether a specific nMSI threshold could serve as a predictive indicator of G4-folding propensity, analogous to the MSI ≥ 0.80 criterion noted for the oncogene promoter sequences in Table 3. Scrutiny of the nMSI values across all four validation datasets (Tables 6–9) revealed a natural and empirically grounded separation between G4-forming and non-G4-forming sequences at nMSI = 0.50. The complete distribution of nMSI values for G4-forming and non-G4-forming sequences across all tables is shown in Figure 4.

**Figure 4.**
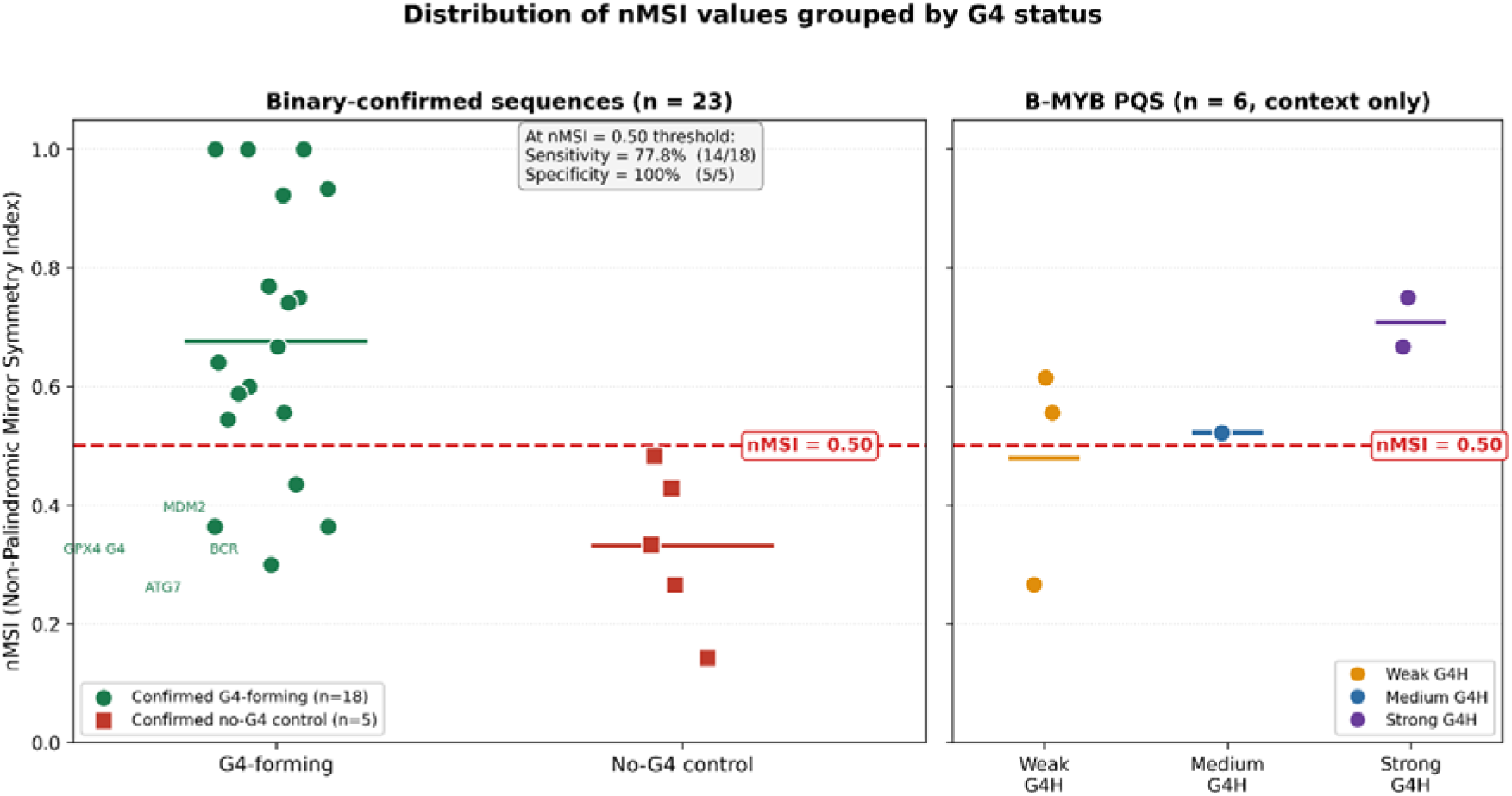
Distribution of Non-Palindromic MSI (nMSI) values grouped by G4 status. The left panel shows the 23 sequences with a binary, experimentally confirmed G4 status—18 confirmed G4-forming sequences (the 14 oncogene-promoter wild-type G4s of Table 6, MYC WT and KRAS SWP of Table 8, and GPX4 G4 and CLIC4 G4 of Table 9) and five confirmed no-G4 controls (the CD-verified G4-abolished mutants MYC MUT, MUT CORE, and MUT MIN of Table 8, and the G4-ChIP-negative controls mutGPX4 and mutCLIC4 of Table 9). Each circle represents a sequence. The red dashed line marks the proposed nMSI = 0.50 threshold, which classifies these binary-confirmed sequences with a sensitivity of 77.8% (14 of 18 confirmed G4-forming sequences correctly identified) and specificity of 100% (all five confirmed no-G4 controls correctly excluded). The right panel shows the six B-MYB promoter PQS (Table 7), which are graded by G4Hunter propensity rather than binary-confirmed and are displayed for context only; their group means illustrates that the nMSI = 0.50 boundary coincides with the transition between weak and at-least-moderate G4-forming propensity.

Among all confirmed no-G4 sequences in the manuscript, comprising the CD-verified G4-abolished mutants MYC MUT, MUT CORE, and MUT MIN (Table 8), and the G4-ChIP-negative controls mutGPX4 and mutCLIC4 (Table 9), the nMSI values span the range 0.143 to 0.483, with the highest observed value being 0.483 for MUT MIN; every confirmed no-G4 control therefore lies below the nMSI = 0.50 boundary. The majority of confirmed G4-forming sequences lie above it, with c-KIT (0.545) being the lowest of the clearly separated cases. A small number of genuinely G4-forming but structurally atypical motifs — the long, loop-rich BCR (0.364), ATG7 (0.300), and MDM2 (0.435) oncogene G4s, and the A-rich-flanked GPX4 G4 (0.364) — fall below the threshold and overlap the control range; these are the loop-architecture and flanking-composition exceptions analyzed below rather than metric failures. Restricting the evaluation to the 23 sequences with a binary, experimentally confirmed G4 status—18 confirmed G4-forming sequences and 5 confirmed no-G4 controls—the nMSI = 0.50 threshold yielded a sensitivity of 77.8% (14 of 18 confirmed G4-forming sequences correctly identified) and a specificity of 100% (all 5 confirmed no-G4 controls correctly excluded). The strand-flipped artifact MYC FLIP and the graded-propensity B-MYB PQS panel (Table 7) were excluded from this binary calculation and considered separately.

A few confirmed G4-forming sequences scored below the nMSI = 0.50 threshold and merited individual consideration. Among the oncogene-promoter panel, the clearest cases are the long, loop-rich ATG7 (nMSI = 0.300) and MDM2 (nMSI = 0.435) motifs, which the present analysis consistently identified as structurally anomalous. The specific loop architectures of these G4s generate non-palindromic trinucleotide pairs that are genuinely and substantially imbalanced, yielding negative Z-scores in the shuffle comparison (Z = −1.202 and −1.557, respectively; Table 5). This is a biologically interpretable result rather than a failure of the metric. Extended asymmetric loop regions impose sequence requirements that are inherently asymmetric at the trinucleotide level, and these cases reveal that nMSI, like MSI, is sensitive to loop architecture. HIF-1α, in contrast, retains an MSI of 0.939 and an nMSI of 0.667, which comfortably exceeded the 0.50 threshold. The only anomaly for HIS-1α was the reversed wild-type versus mutant comparison (Table 6), which arose from a coincidentally palindrome-rich scrambled mutant rather than from low wild-type symmetry. A further below-threshold example is GPX4 G4 (nMSI = 0.364, Table 9), a G4-ChIP-confirmed sequence who’s depressed nMSI arises from its five-nucleotide A-rich 5′ prefix (AAAAC…) preceding the G4-forming core. These A-rich flanking residues introduce asymmetric non-palindromic trinucleotides unrelated to the G-quartet-forming region, diluting the nMSI below the threshold, despite genuine G4 formation being confirmed in living cells. The original MSI (0.857) is less susceptible to this effect because palindromic GGG trinucleotides within the G4 core dominate the MSI denominator and partially offset flank-induced asymmetry. These two exceptions suggest that nMSI ≥ 0.50 should be interpreted in conjunction with the MSI. Sequences that satisfy MSI ≥ 0.80 but fall below nMSI = 0.50 warrant further scrutiny of their topology or flanking composition before being excluded as non-G4-forming.

The single apparent exception on the no-G4 side is MYC FLIP (nMSI = 0.720, Table 8), which scores above the threshold despite being confirmed by CD spectroscopy to lack G4 formation. As noted in the discussion of Table 8, MYC FLIP is the reverse-complement of MYC WT repositioned on the template strand, and its trinucleotide composition is identical to that of MYC WT. Because nMSI, like MSI, is a composition-based metric, it necessarily assigns the same score to any two sequences sharing identical trinucleotide frequencies, regardless of strand identity. Therefore, MYC FLIP is not a biological false positive but a known and acknowledged compositional artifact, a limitation shared by all composition-based symmetry indices and explicitly discussed in the context of MSI earlier in this work.

The complete distribution of MSI values for G4-forming and non-G4-forming sequences across all tables is shown in Figure S4. A careful reading of the distributions of MSI and nMSI over the sequences studies indicates that the nMSI ≥ 0.50 threshold offers several practical advantages over the MSI ≥ 0.80 threshold. First, and most critically, an MSI ≥ 0.80 failed to correctly classify several sequences in the external validation datasets. CLIC4 G4, a hybrid G4 confirmed by G4-ChIP in living cells (Table 9), has an MSI of only 0.789, which falls below the 0.80 threshold and would lead to its exclusion as a non-G4-forming candidate despite direct cellular evidence of G4 folding. In contrast, nMSI correctly identified this sequence with a value of 0.556, which was comfortably above 0.50. Conversely, MUT CORE and MUT MIN — sequences in which G4 formation has been abolished, as confirmed by CD spectroscopy — have MSI values of 0.750 and 0.762, respectively, placing them disconcertingly close to the 0.80 boundary and illustrating the compressed dynamic range of the original MSI for this class of sequences. Their nMSI values of 0.429 and 0.483 fell below the 0.50 threshold, providing a cleaner and more confident separation. Second, the nMSI threshold aligns well with the graded G4-propensity series in Table 7. Across the B-MYB promoter PQS panel, strong G4 sequences (G4H score ∼2.9) showed a mean nMSI of 0.709, the medium-strength candidate scored 0.522, and weak G4 sequences averaged 0.479. The nMSI = 0.50 boundary thus falls precisely at the transition between weak and at least moderate G4-forming propensity, lending the threshold an interpretable biological basis beyond the binary classification itself. Third, the enhanced dynamic range of nMSI, a consequence of excluding the palindromic trinucleotide ceiling that compresses MSI values toward 1.0 in G-rich sequences, means that nMSI discriminates more finely within the G4-forming class itself, with values spanning 0.300–1.000 across the oncogene panel compared with the narrower effective range of 0.650–1.000 observed for the same 14 oncogene WT sequences in MSI. This expanded spread is precisely what enables a lower absolute threshold (0.50 vs. 0.80) to achieve comparable or superior specificity, with only a modest reduction in sensitivity.

Taken together, these observations indicate that an nMSI ≥ 0.50 criterion is a useful and data-supported adjunct to the existing MSI ≥ 0.80 threshold for identifying sequences with G4-folding propensity. The two metrics are most appropriately used in combination: MSI ≥ 0.80 captures the strong palindromic-symmetry signal characteristic of the most G-rich, short oncogene promoter G4s, while nMSI ≥ 0.50 provides complementary discrimination that is more sensitive to asymmetric non-palindromic pair balance, more robust across diverse sequence contexts including non-G4-rich flanking regions, and less susceptible to the ceiling effect that limits MSI’s discriminatory power. Sequences satisfying both criteria can be considered high-confidence G4-forming candidates; sequences satisfying nMSI ≥ 0.50 but not MSI ≥ 0.80, or vice versa, warrant follow-up analysis, accounting for topology and sequence composition.

The MSI and nMSI as presently formulated face the following limitations: (i)□with the exception of KRAS (p = 0.042 on MSI; p = 0.0349 on nMSI), the per-sequence shuffle p-values do not reach significance, reflecting three deeper compounding factors: (a)□the short sequence lengths of these motifs yield few trinucleotide observations; (b)□extreme G-richness causes palindromic trinucleotides, especially GGG, to constitute a large fraction of the original MSI denominator while contributing zero to the numerator, pushing both WT and shuffled MSI toward the same ceiling, which is a confound addressed by nMSI (Eq. □2); (c)□the constrained mononucleotide-preserving shuffle space for such G-rich sequences limits the distributional width. A global combined test (Stouffer’s method) applied to nMSI Z-scores across all 14 sequences yielded Z□_combined_□=□1.197, p□=□0.116, which represents a non-significant collective trend rather than definitive per-sequence evidence; the strongest statistical support instead comes from the wild-type versus mutant sign test (p = 0.0065). (ii)□Two motifs (BCL2 and HIF-1α) show a reversed wild-type-versus-mutant relationship attributable to scrambled-sequence mutant controls that coincidentally introduce more balanced trinucleotide compositions; therefore, gene-specific, experimentally validated mutants should be used wherever available. (iii)□The mechanistic link between mirror symmetry and G4 folding remains a hypothesis requiring experimental validation.

## Conclusion

The results and analysis presented in this study indicate that the insertion of nucleotides in both strands of DNA simultaneously and the maintenance of quadruplets represent an evolutionary forcing mechanism that gives rise to genomic organization at all scales[3], [8]. This forcing mechanism operates at the local scale in functional elements, such as G4-forming sequences, to generate selective pressure to maintain trinucleotide symmetries. This perspective justifies several apparently intriguing observations: (1) the observation of CSPR in the face of local violations in coding regions; (2) the observation of functional G4s to be symmetric in the face of codon-bias constraints; and (3) the selective preservation of G4 sequences over evolutionary time. In addition, the proposed framework implies that symmetry breaks in certain genomic contexts (e.g., disease-related mutations disrupting the trinucleotide symmetry in G4 motifs) can have functional effects in addition to merely disrupting the structure. Such symmetry breaks can be viewed as a violation of the general law of evolution, with an impact on gene regulation and disease pathophysiology[59]. An example of such a principle is the conserved G4 motif in the first exon of the MTOR gene: although this sequence is in the coding DNA where codon-bias-induced asymmetries can occur, it still exhibits high trinucleotide symmetry. This increase is indicative of natural selection that maximizes the creation of G4 structures. Experimentally validated G4s in oncogenes (c-MYC, BCL2, VEGF, and KRAS) with comparative analysis showed that the mirror and reverse-complement trinucleotide symmetries converge around functionally important G4s and indicate that symmetry is central to G4 biology. Mirror symmetry may pre-organize the single strand for G4 folding by reducing the entropic cost of loop formation and facilitating favorable loop–core interactions. To quantify this organizational property, we formulated two complementary sequence-based descriptors, the mirror symmetry index (MSI) and its non-palindromic variant (nMSI), which act as prioritization filters that complement existing G-tract-based prediction tools. Applied across 14 oncogene promoter G4s, most wild-type sequences scored MSI ≥ 0.80 (mean 0.884), and the KRAS promoter G4 reached individual significance against its mononucleotide-preserving null distribution (p = 0.042). The most decisive evidence emerged from comparing each wild-type G4 against its experimentally G4-abolished mutant: the wild-type scored higher in 12 of 14 paired comparisons (sign test, p = 0.0065; mean ΔMSI = +0.089, mean ΔnMSI = +0.192), a single global test that established a robust directional association between mirror symmetry and G4-forming capacity. The two reversals, BCL2 and HIF-1α, are attributable to scrambled-sequence mutant controls that coincidentally adopt more balanced trinucleotide compositions rather than any failure of the index. This directional trend was reproduced across three independent, recently published datasets spanning CRISPR-edited, graded-propensity, and cellularly validated G4-ChIP sequences. The nMSI was introduced to overcome a systematic confound whereby palindromic trinucleotides, chiefly GGG, dominate the MSI denominator in these G-rich sequences while contributing no asymmetry information, compressing both wild-type and shuffled values toward a common ceiling of approximately 1.0. By restricting the index to asymmetric trinucleotide pairs, nMSI sharpens the discrimination between G4-forming and non-G4-forming sequences, even though the per-sequence shuffle trend across the panel remained non-significant (Stouffer Z = 1.197, p = 0.116). We further identified an empirically grounded nMSI ≥ 0.50 threshold that separates confirmed G4-forming from non-G4-forming sequences across the validation datasets with 77.8% sensitivity and 100% specificity across the 23 binary-confirmed sequences, with every confirmed no-G4 control lying below the boundary, and the boundary coinciding with the transition between weak and moderate G4 propensity in a graded series. The two indices are best used in combination: MSI ≥ 0.80 captures the strong palindromic-symmetry signal of the most G-rich, short promoter G4s, whereas nMSI ≥ 0.50 provides finer, composition-robust discrimination that correctly rescues cellularly confirmed sequences such as CLIC4, which the MSI threshold alone would misclassify. The few sequences that deviate from these thresholds, including the anti-parallel/mixed HIF-1α G4 and the A-rich-flanked GPX4 G4, are biologically interpretable consequences of topology or flanking composition rather than failures of the metrics, reinforcing that mirror symmetry is sensitive to G4 architecture. While the MTOR gene served as the primary model for evolutionary conservation analysis, the structural and functional diversity of the validation sequences, spanning parallel, anti-parallel, and mixed G4 topologies and multiple independent experimental modalities, supports the generalizability of both indices. We acknowledge that the per-sequence shuffle tests are individually underpowered owing to the short lengths and extreme G-richness of these motifs and that the mechanistic coupling between mirror symmetry and G4 folding remains a hypothesis awaiting direct experimental testing. The extension of the MSI/nMSI framework to RNA G4s, telomeric repeats, and genome-wide G4-ChIP datasets represents an important direction for future studies. Finally, this study places the Natural Symmetry Law as a general evolutionary force, which limits genomic-scale organization, encompassing the balance of whole-genome nucleotides and the structure of functional regulatory elements.

## Author Information

### Author Contributions

Arjun Arya: Conceptualization, Investigation, Methodology, Data curation, Validation, Formal Analysis, Visualization, Writing – original draft, Writing – review & editing. Bhaskar Datta: Conceptualization, Methodology, Validation, Writing – original draft, Writing – review and editing, Funding acquisition, Project administration, Resources, Supervision.

## Acknowledgements

B.D. gratefully acknowledges financial support for this work from the Gujarat State Biotechnology Mission (GSBTM) (project no. GSBTM/JD(R&D)/626/22-23/00006262. All illustrations were created using Biorender.com.

## Notes

### Competing Interest Statement

The authors have declared no competing interest.

