## Supplementary Information for "Trinucleotide Distribution, Symmetry Elements and Formulation of Mirror Symmetry Index for G4 Motifs"

* Corresponding author.

Table S1. Nucleotide distribution patterns of MTOR homologs.

| **Organism** | **GC-content** | **AT-content** | **A%** | **C%** | **G%** | **T%** |
| --- | --- | --- | --- | --- | --- | --- |
| **Human** | 46.76 | 53.24 | 25.36 | 22.88 | 23.88 | 27.88 |
| **Chimp** | 46.54 | 53.46 | 25.12 | 22.75 | 23.79 | 23.79 |
| **Monkey** | 46.12 | 53.88 | 25.33 | 22.47 | 23.65 | 28.55 |
| **Gorilla** | 46.51 | 53.49 | 25.1 | 22.71 | 23.8 | 28.39 |
| **Pig** | 50.57 | 49.43 | 22.13 | 25.03 | 25.54 | 27.3 |
| **Dog** | 45.9 | 54.1 | 24.99 | 22.43 | 23.47 | 29.12 |
| **Mouse** | 49.09 | 50.91 | 22.85 | 23.74 | 25.35 | 28.06 |
| **Rat** | 48.76 | 51.24 | 23.37 | 23.88 | 24.88 | 27.86 |
| **Cattle** | 45.13 | 54.87 | 24.74 | 22.37 | 22.75 | 30.13 |

**Table S2.** Classification of trinucleotides showing purine-pyrimidine symmetry and Watson-Crick base pairing patterns in G-quadruplex-forming sequences. Trinucleotides are organized into A+T-rich (Group I) and C+G-rich (Group II) categories, further subdivided into symmetry-based subgroups (a, b, c). Each trinucleotide is presented in four Watson-Crick transformation forms: direct (D), reverse complement (RC), complement (C), and reverse (R). Binary notation (0 = purine, 1 = pyrimidine) beneath each sequence reveals purine-pyrimidine symmetry patterns. Asterisks (*) indicate self-complementary trinucleotides. This systematic organization demonstrates that trinucleotide composition follows predictable mathematical principles and may represent evolutionarily conserved structural features in G-quadruplex sequences.

| **A+T Rich Group (I)** | | | | | **C+G Rich Group (II)** | | | | |
| --- | --- | --- | --- | --- | --- | --- | --- | --- | --- |
| **D** | **RC(D)** | **C(D)** | **R(D)** | **Subgroup** | **D** | **RC(D)** | **C(D)** | **R(D)** | **Subgroup** |
| ATG | CAT | TAC | GTA | Ia | CGT | ACG | GCA | TGC | IIa |
| 010 | 101 | 101 | 010 |  | 101 | 010 | 010 | 101 |  |
| TGA | TCA | ACT | AGT |  | GTC | GAC | CAG | CTG |  |
| 100 | 110 | 011 | 001 |  | 011 | 001 | 100 | 110 |  |
| TAG | CTA | ATC | GAT |  | GCT | AGC | CGA | TCG |  |
| 100 | 110 | 011 | 001 |  | 011 | 001 | 100 | 110 |  |
| TAA | TTA | ATT | AAT | Ib | GCC | GGC | CGG | CCG | IIb |
| 100 | 110 | 011 | 001 |  | 011 | 001 | 100 | 110 |  |
| AAC | GTT | TTG | CAA |  | CCA | TGG | GGT | ACC |  |
| 001 | 011 | 110 | 100 |  | 110 | 100 | 001 | 011 |  |
| AAG | CTT | TTC | GAA |  | CCT | AGG | GGA | TCC |  |
| 000 | 111 | 111 | 000 |  | 111 | 000 | 000 | 111 |  |
| ATA | TAT | TAT* | ATA* | Ic | CGC | GCG | GCG* | CGC* | IIc |
| 010 | 101 | 101 | 010 |  | 101 | 010 | 010 | 101 |  |
| ACA | TGT | TGT* | ACA* |  | CAC | GTG | GTG* | CAC* |  |
| 010 | 101 | 101 | 010 |  | 101 | 010 | 010 | 101 |  |
| AGA | TCT | TCT* | AGA* |  | CTC | GAG | GAG* | CTC* |  |
| 000 | 111 | 111 | 000 |  | 111 | 000 | 000 | 111 |  |
| AAA | TTT | TTT* | AAA* |  | CCC | GGG | GGG* | CCC* |  |
| 000 | 111 | 111 | 000 |  | 111 | 000 | 000 | 111 |  |

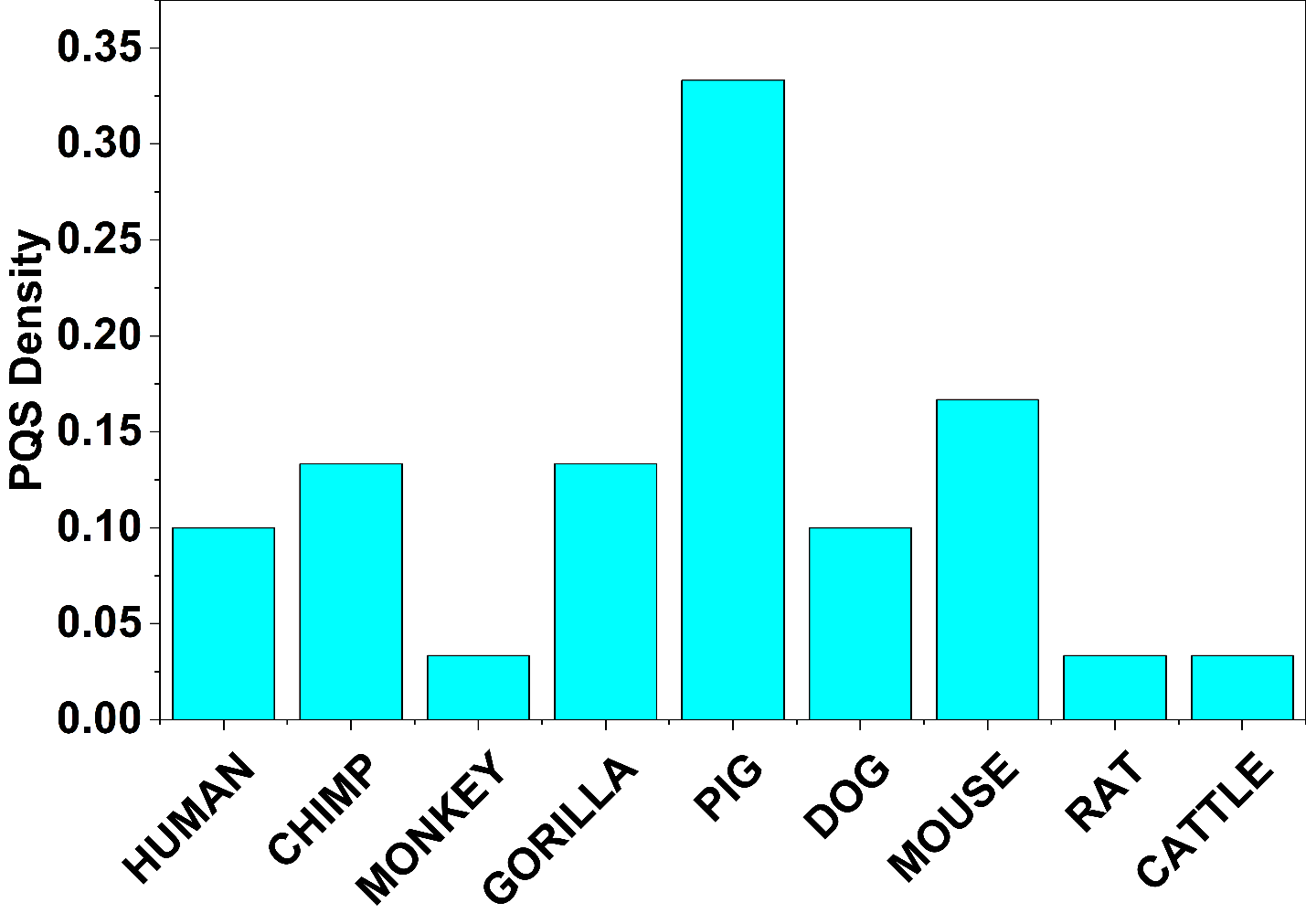

**Figure S1.** Comparative predicted PQS densities in the MTOR upstream region across organisms.

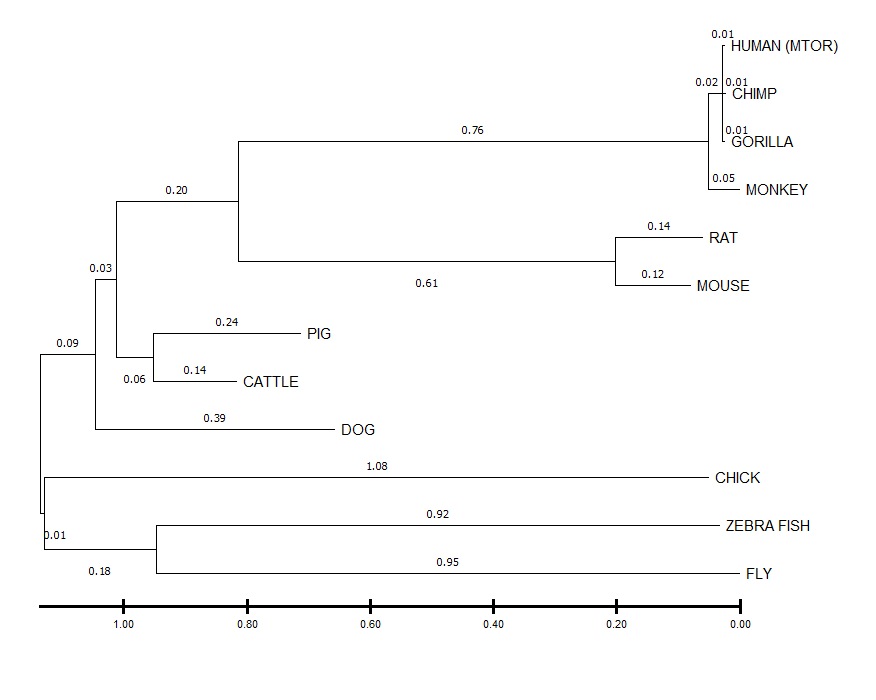

**Figure S2.** Unrooted phylogenetic tree showing evolutionary relationships based on mTOR G4 sequence divergence. Branch lengths indicate evolutionary distance (substitutions per site), with values labelled at major nodes.

(b)

(a)

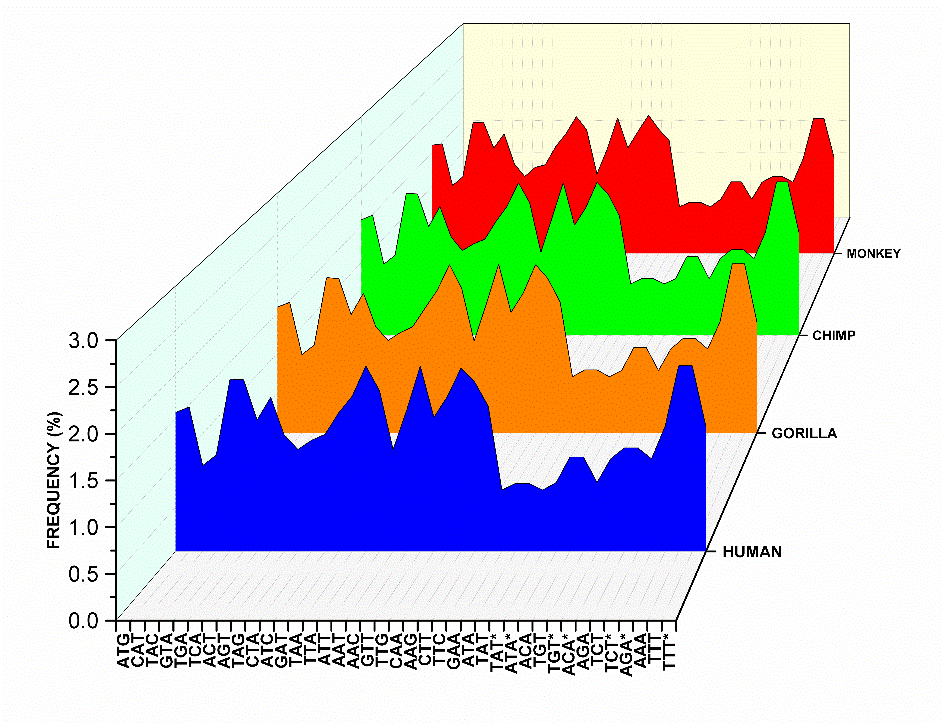

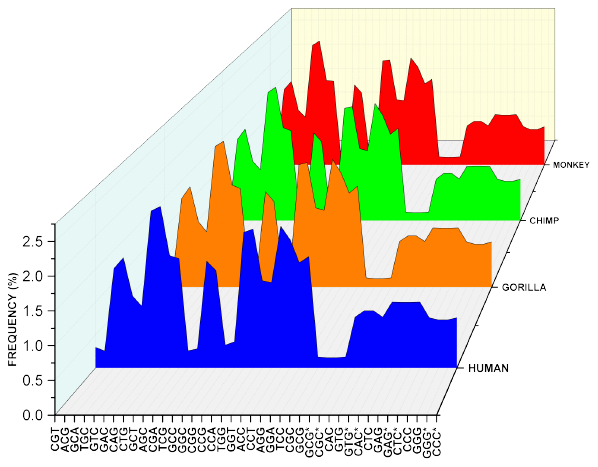

**Figure S3.** Frequency distribution of quadruplets on the MTOR gene across primate species. Area plots showing frequency (%) of (a) A+T-rich quadruplets and (b) C+G-rich quadruplets of the MTOR gene in human (blue), gorilla (orange), chimp (green), and monkey (red). The analysis reveals species-specific variation in the frequencies of AT-rich and GC-rich trinucleotide motifs across the MTOR gene.

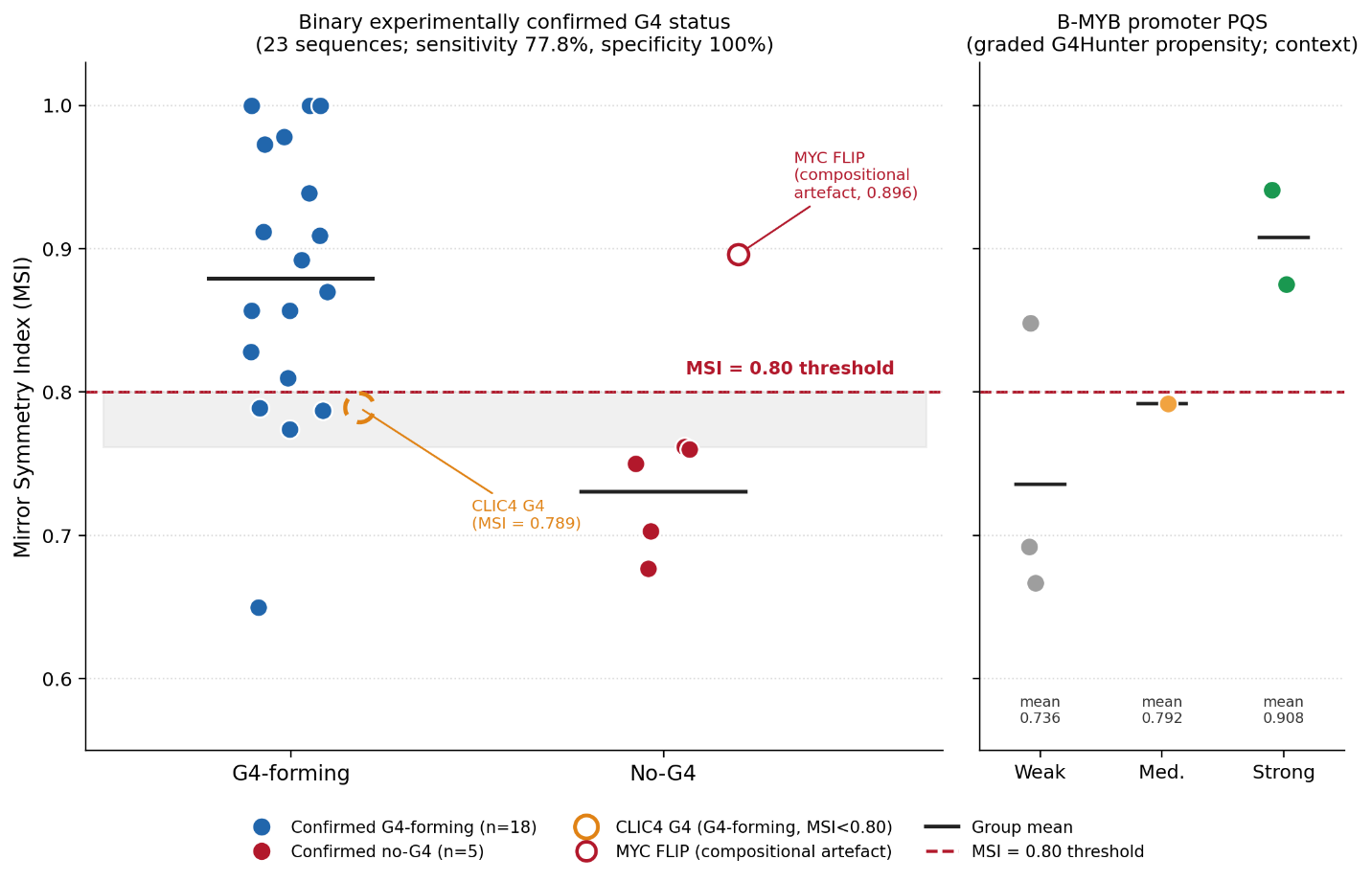

**Figure S4.** Distribution of Mirror Symmetry Index (MSI) values grouped by experimentally determined G4 status, from Tables 4, 6, 7, 8 and 9. The left panel shows the 23 binary-confirmed sequences (18 confirmed G4-forming, 5 confirmed no-G4). The red dashed line marks the MSI = 0.80 threshold, which classifies these sequences with 77.8% sensitivity (14/18) and 100% specificity (5/5); the shaded band spans the natural gap between the highest no-G4 control (MUT MIN, 0.762) and the threshold. CLIC4 G4 (amber ring) is a G4-ChIP-confirmed sequence that falls just below 0.80 (MSI = 0.789) but is correctly recovered by nMSI ≥ 0.50 (nMSI = 0.556). MYC FLIP (hollow red) exceeds the threshold but is an acknowledged compositional artefact. The right panel shows the six graded B-MYB PQS (Table 7) for context.
